# Human monoclonal antibodies reveal subdominant gonococcal and meningococcal cross-protective antigens

**DOI:** 10.1101/2023.12.07.570438

**Authors:** Marco Troisi, Monica Fabbrini, Samuele Stazzoni, Viola Viviani, Filippo Carboni, Valentina Abbiento, Lucia Eleonora Fontana, Sara Tomei, Martina Audagnotto, Laura Santini, Angela Spagnuolo, Giada Antonelli, Ida Paciello, Fabiola Vacca, Dario Cardamone, Eleonora Marini, Pardis Mokhtary, Francesca Finetti, Fabiola Giusti, Margherita Bodini, Giulia Torricelli, Chiara Limongi, Mariangela Del Vecchio, Sara Favaron, Simona Tavarini, Chiara Sammicheli, Alessandro Rossi, Andrea Paola Mandelli, Pietro Fortini, Carla Caffarelli, Stefano Gonnelli, Ranuccio Nuti, Cosima T. Baldari, Claudia Sala, Aldo Tagliabue, Silvana Savino, Brunella Brunelli, Nathalie Norais, Elisabetta Frigimelica, Monia Bardelli, Mariagrazia Pizza, Immaculada Margarit, Isabel Delany, Oretta Finco, Emanuele Andreano, Rino Rappuoli

## Abstract

Gonococcus (Gc), a bacterium resistant to most antibiotics causing more than 80 million cases of gonorrhea annually, is a WHO high priority pathogen. Recently, vaccine development prospects were boosted by reports that licensed meningococcus serogroup B (MenB) vaccines provided partial protection against Gc infection. To determine antigens responsible for cross-protection, memory B cells from 4CMenB vaccinated volunteers were single-cell sorted to identify antibodies that kill Gc in a bactericidal assay. Nine different antibodies, all deriving from the IGHV4-34 germline carrying unusually long HCDR3s, recognized the PorB protein, four recognized the lipooligosaccharide (LOS), and four unknown antigens. One of the PorB antibodies, tested in vivo, provided protection from Gc infection. The identification of PorB and LOS as key antigens of gonococcal and meningococcal immunity provides a mechanistic explanation of the cross-protection observed in the clinic and shows that isolating human monoclonal antibodies from vaccinees can be instrumental for bacterial antigen discovery.

## INTRODUCTION

*Neisseria gonorrhoeae* (Gonococcus; Gc), the causative agent of gonorrhea, has been a persistent public health problem for centuries (Hill et al., 2016). Today, gonorrhea is the second most common sexually transmitted disease, causing 710,000 infections yearly in the United States and more than 80 million cases globally (St Cyr et al., 2020; Unemo et al., 2021). Gonococcal infection can lead to pelvic inflammatory disease, infertility, and ectopic pregnancies (Lenz and Dillard, 2018). Moreover, the impact of gonorrhea on human health is amplified by its role in increasing both transmission and susceptibility to the human immunodeficiency virus 1 (HIV-1) (Jarvis and Chang, 2012). Since the 1940s the disease has been treated with antibiotics, however, over time the bacterium has acquired resistance to sulphonamides, penicillins, fluoroquinolones and today it is only susceptible to third generation cephalosporins (Unemo and Shafer, 2014). The recent isolation of strains resistant even to cephalosporins (Młynarczyk-Bonikowska et al., 2020; Unemo and Shafer, 2014) raised the alarm of the World Health Organization (WHO) that, concerned about the bacterium becoming untreatable, concluded that the development of new antimicrobials and Gc vaccines is imperative (Goire et al., 2014; Unemo et al., 2021; Unemo and Shafer, 2014). The search for vaccines against Gc has been largely unsuccessful for many decades (Russell et al., 2019). Although in 1978 a human challenge trial demonstrated protection against infection using a pilus purified from the challenge strain as a vaccine, a large-scale phase 3 trial involving 3,250 volunteers failed to show any efficacy, most likely because of the large antigenic variability of the antigen used in this vaccine (Boslego et al., 1991; Gottlieb et al., 2020; Greenberg, 1975; Maurakis and Cornelissen, 2022; Russell et al., 2019; Tramont, 1989). After decades of failures in developing gonococcal vaccines, the recent observation of partial efficacy against Gc infection of *Neisseria meningitidis* serogroup B (MenB) Outer Membrane Vesicle (OMV)-based vaccines has revived the hopes for gonococcal vaccine research (Paynter et al., 2019; Semchenko et al., 2019). Briefly, a retrospective case-control study conducted in New Zealand following mass vaccination campaign with an OMV based MenB vaccine (MeNZB) (Paynter et al., 2019) and three observational studies conducted in the U.S. and Australia (Abara et al., 2022; Bruxvoort et al., 2023; Wang et al., 2022) reported 30-46% protection against Gc infection and confirmed observations that had been made in the past in Canada (Longtin et al., 2017), Cuba (Ochoa-Azze, 2018) and Norway (Whelan et al., 2016). These observations provided proof of concept that a Gc vaccine is feasible and suggested that MenB OMV-based vaccines contained antigens that might induce a cross-reactive immunity against Gc. The hypothesis is supported by the fact that MenB and Gc share between 80 and 90% of genome identity (Tinsley and Nassif, 1996) and their lipooligosaccharide (LOS) antigens share partial similarity (Mandrell et al., 1988). Since the PorA immunodominant protective antigen of the meningococcal OMV vaccine is not expressed by Gc (Unemo et al., 2005), it has been hypothesized that other MenB subdominant antigens might be responsible for the observed cross-protection (Semchenko et al., 2019). A comparative genomic analysis on approximately 1,000 Gc isolates collected in the U.S. and MenB reference strains allowed the identification of 57 Outer Membrane Proteins (OMPs) with high homology between the two Neisseria species (Marjuki et al., 2019). Approximately 42% of common OMPs have been identified in the OMV component of the 4CMenB vaccine licensed against MenB (Ferrari et al., 2006; Holst et al., 2013; Tani et al., 2014; Vipond et al., 2006). These included BamA, NspA, MtrE and MetQ, exhibiting between 91% and 100% amino acid sequence similarity, and PorB, RmpM, PilQ, OpcA, FetA, Omp85 (BamA) and LbpA, which were shown to be consistently present in different OMV lots of the 4CMenB vaccine (Tani et al., 2014). In addition, the 4CMenB vaccine contained also the NHBA protein which is moderately homologous between the two *Neisseria* species (Marjuki 2019). To understand which MenB antigens could contribute to the cross-protection observed against Gc, we immunized healthy volunteers with the 4CMenB vaccine, collected their peripheral blood cells (PBMCs), single cell sorted the memory B cells (MBCs) and selected the B cells producing monoclonal antibodies (mAbs) recognizing the meningococcal OMV and killing gonococcus *in vitro* (**Fig. 1**). This approach allowed the identification of PorB and LOS as key antigens for cross-protection against gonococcus and meningococcus and provided a paradigm that could be used for antigen discovery for other antibiotic resistant bacteria.

**Fig. 1.**
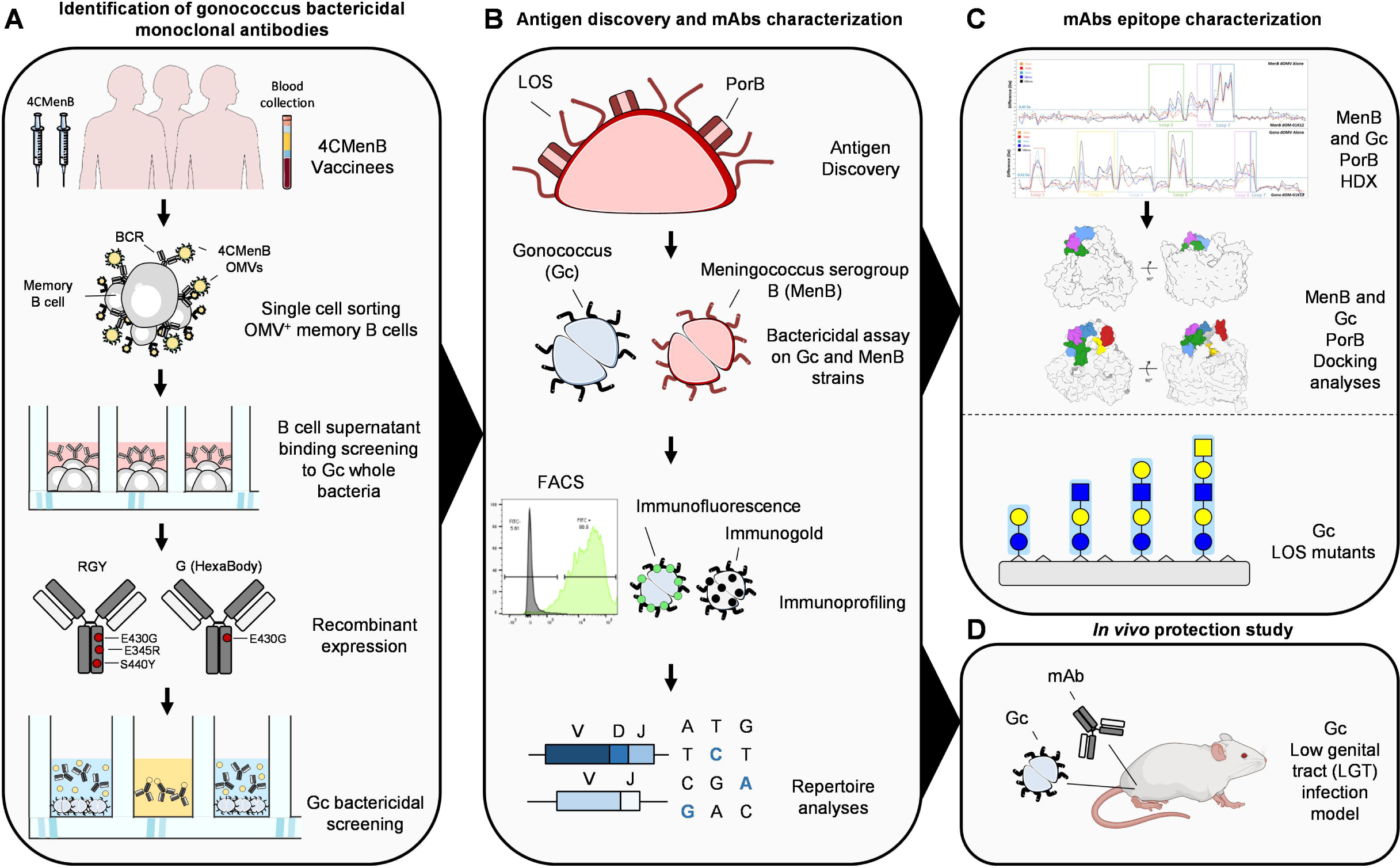
Workflow for Gc bactericidal mAbs identification and characterization. The overall scheme shows four different steps to identify and characterize bactericidal mAbs. (**A**) The first steps consist in the enrolment of 4CMenB vaccinees (n=3) from which blood was collected and PBMCs isolated. 4CMenB OMVs^+^ memory B cells were single cell sorted (n=3,080) and after 2 weeks of incubation B cell supernatant were screened for their binding specificity against different Gc strains. Once identified, Gc binding mAbs (n=390) were recombinantly expressed in RGY and G (HexaBody) scaffolds to evaluate their bactericidal activity to Gc. (**B**) The second step starts with the characterization of identified functional mAbs in the HexaBody scaffold (n=17). Firstly, antibodies were tested to identify their cognate antigen. Next, mAbs were evaluated for their ability to kill different Gc and MenB strains, and for their binding profiles. Finally, the heavy and light chain sequences of mAbs were recovered and repertoire analysis performed. (**C**) Selected anti-PorB and all anti-LOS mAbs were further characterized to identify their targeted epitope. (**D**) In addition, the most promising anti-PorB mAb was tested *in vivo* to evaluate its efficacy in preventing Gc infection.

## RESULTS

### Identification of bactericidal mAbs (b-mAbs) against Gc

To isolate MBCs specific for MenB OMVs contained in the 4CMenB formulation, 3 volunteers were vaccinated with the 4CMenB vaccine and their PBMCs were collected 28 days after receiving the booster dose (**Fig. S1A**). PBMCs were stained with antibodies for CD19, CD27, IgD, IgM and MenB OMVs labelled with Alexa488, with the aim to identify class-switched memory B cells (MBCs; CD19^+^CD27^+^IgD^-^IgM^-^) OMV-binders. The gating strategy applied to perform the single cell sorting is described in **Fig. S1B**. Using this approach 3,080 OMV^+^ MBCs were sorted and incubated for 2 weeks to allow natural production of immunoglobulins as previously described (Andreano et al., 2021). The percentage of class-switched MBCs detected in the three donors was 23.2, 16.9 and 43.1% for subjects 7, 8 and 9, respectively, and the percentage of OMV^+^ MBCs ranged from 2.6 to 12.6% (**Fig. S1C**). To identify OMV^+^ MBCs that could cross-bind Gc, we performed a whole bacterial cell enzyme-linked immunosorbent assay (ELISA) with the Gc strains FA1090 and F62 (Hobbs et al., 2011), and the recently described low-passaged clinical isolate BG27 (Manca et al., 2023). This latter strain expresses the same PorB.1B variant of FA1090 and presents a polyphosphate (polyP) pseudo-capsule that has been shown to confer high serum resistance. From the 3,080 OMV^+^ MBCs sorted, a panel of 390 (12.7%) antibodies recognized at least one of the tested Gc strains in ELISA (**Fig. 2A**). The variable regions of the mAbs heavy (VH) and light (VL) chains were cloned into appropriate vectors for mAb expression as previously described (Andreano et al., 2021). All 390 Gc-binding mAbs were expressed in small scale (1 mL) through transcriptionally active polymerase chain reaction (TAP) and screened by resazurin-based high-throughput antibody bactericidal assay (R-ABA) (Stazzoni et al., 2023). For this assay, mAbs were incubated with Gc strains in presence of baby rabbit complement (10-20%) and bacterial viability was measured through the use of resazurin. To maximize the sensitivity of the bactericidal assay, the antibodies were initially engineered using a fragment crystallizable (Fc) region carrying the RGY mutations (E345R, E430G, S440Y) which promote antibody hexamerization and enhance C1q deposition on the bacterial surface (de Jong et al., 2016; McIntosh et al., 2015). Among the 390 tested mAbs, 36 (9.2%) were found to be bactericidal against FA1090 (**Fig. 2B**). However, while the RGY antibody scaffold is useful for screening, it cannot be used clinically because of non-specific antibody hexamerization and complement binding in solution even in the absence of the target antigen (de Jong et al., 2016). Therefore, the subsequent work was performed using a scaffold carrying only the G (E430G) mutation, known as HexaBody (de Jong et al., 2016). This scaffold still enables improved complement deposition compared to natural IgG1, while preventing target-independent hexamerization and C1q binding in solution. In addition, the HexaBody scaffold is currently being used in different trials supporting its selection for clinical development (De Goeij et al., 2019; Oostindie et al., 2020). Seventeen out of 36 (47.2%) b-mAbs expressed as HexaBody retained their bactericidal activity against strain FA1090 with potency values (50% inhibitory concentration; IC*_50_*) ranging from 0.05 to ∼150 µg/mL (**Fig. 2C and S2**).

**Fig. 2.**
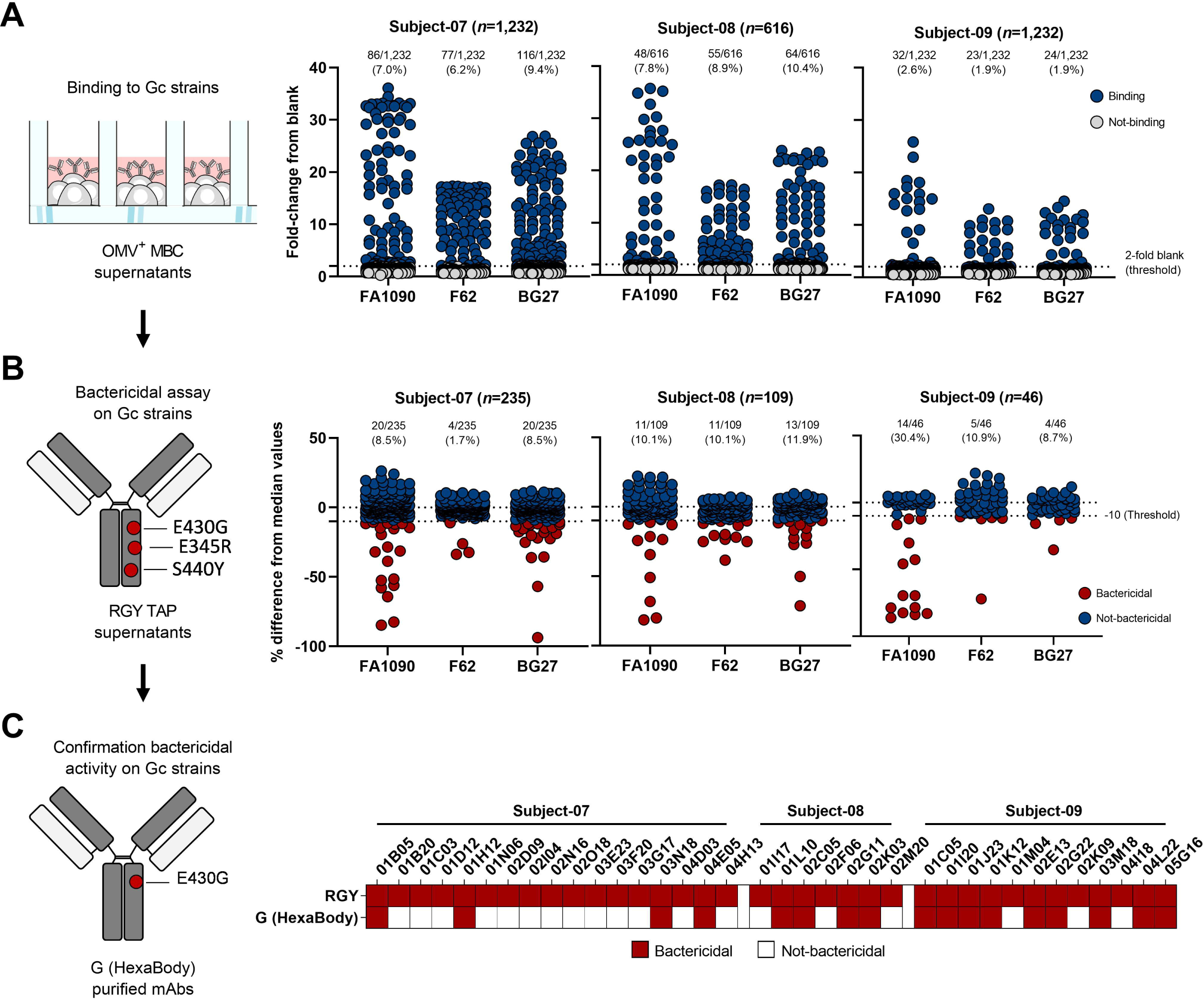
Identification of Gc bactericidal mAbs. (**A**) The graphs show 4CMenB OMV^+^ mAbs binding activity against Gc strains FA1090, F62 and BG27. The threshold of positivity was set as two times the value of the blank (dotted line). The dark blue and gray dots represent binding and non-binding mAbs respectively. The number and percentage of mAbs tested per donor are denoted on each graph. (**B**) Dot charts show the bactericidal activity of Gc-binding mAbs recombinantly expressed in the RGY scaffold. The threshold of positivity was set as 10% signal reduction from medium value (dotted lines). The dark red and dark blue dots represent bactericidal and not-bactericidal mAbs respectively. The number and percentage of mAbs tested per donor are denoted on each graph. (**C**) The heatmap shows the bactericidal activity against Gc FA1090 strain of mAbs expressed in the G (HexaBody) scaffold. Dark red and white boxes represent bactericidal and non-bactericidal mAbs respectively.

### Antigen identification and functional mAb characterization on Gc and MenB strains

To identify the antigens recognized by the selected mAbs, we performed immunoblot assays using MenB lysed OMVs or purified LOS. In addition, we used microarrays containing 12 recombinant MenB proteins and 26 recombinant *E. coli* Generalized Modules for Membrane Antigens (GMMAs) expressing MenB OMPs (Viviani et al., 2023). Four of the total 17 b-mAbs (01B05, 03N18, 04E05 and 02G11) recognized purified MenB LOS by immunoblot (**Fig. 3A**). Seven b-mAbs tested by immunoblotting on MenB OMVs recognized a band co-migrating with the Porin B (PorB) protein (**Fig. 3B**). The signal of band intensity for these mAbs ranged from high (01J23, 01K12 and 03M18) to weak (01H12, 02C05, 02K03 and 04L22) (**Fig. 3B**). The recognition of PorB by all 7 mAbs, including the weakly reactive ones, was confirmed through protein microarray analysis and binding to purified recombinant MenB PorB and *E. coli* GMMAs expressing MenB PorB (**Fig. 3C**). Moreover, two additional anti-PorB antibodies (01C05 and 02G22) were identified through binding of PorB on the microarray (**Fig. 3C**). Therefore, our results revealed that a total of 9 (53.0%) b-mAbs targeted the PorB antigen and 4 (23.5%) recognized LOS. For the remaining 4 (23.5%) b-mAbs (01L10, 01I20, 02E13 and 05G16) we were unable to identify the target antigen neither by Western blotting nor by protein microarray. To confirm the bactericidal activity of selected identified b-mAbs, antibodies were tested against the Gc strain FA1090 through classical serum bactericidal assay (**Fig. 3D, left panel**). All antibodies were able to kill Gc FA1090 (**Fig. 3D, left panel, Table S1**). The 17 mAbs killing the FA1090 strain were evaluated for phagocytic activity through the visual opsonophagocytosis assay (vOPA). This approach is based on high-content image analysis to count the number of bacteria internalized by macrophage-like THP-1 cells in the presence of b-mAbs (**Fig. 3D, middle panel**). With the exception of anti-PorB mAb 04L22, all antibodies tested showed opsonophagocytic activity, and 4 anti-PorB antibodies, 01J23, 01K12, 01C05 and 02G22, were found to be the most potent in promoting bacterial internalization by THP-1 macrophage-like cells. Finally, we evaluated through microarray analysis the ability of our Gc b-mAbs to cross-react with additional eighteen low-passaged BG clinical strains (**Fig. S3**). Overall, broad cross-reactivity was shown by all b-mAbs with the exception of 02C05 (anti-PorB), 01L10 (unknown target), 03M18 (anti-PorB) and 02E13 (unknown target) which recognized 4, 3, 0 and 0 BG strains respectively. Distinct binding profiles were observed between anti-LOS and anti-PorB b-mAbs (**Fig. S3**). In fact, anti-LOS were the only antibodies able to bind both BG1 and BG11. Conversely, almost all anti-PorB b-mAbs were able to bind BG10, BG19, BG21, BG23, BG24 and BG29 which were not bound by anti-LOS antibodies. Antibodies targeting unknown antigens showed lower cross-reactivity to BG strains, with the exception of 05G16 which presented a binding profile similar to anti-PorB b-mAbs (**Fig. S3**). Since Gc b-mAbs isolated in this study were elicited by the MenB 4CMenB vaccine, we also evaluated the bactericidal activity of 17 antibodies against a panel of ten strains representative of the genetic diversity of MenB (**Fig. 3D, right panel**). The panel includes the strains used to produce the Norwegian and Cuban MenB OMV-based vaccines (H44-76 and CU385), the reference strains for NadA and NHBA 4CMenB antigens (5/99, NGH38) (Donnelly et al., 2010), the New Zealand epidemic strain (NZ98/254) from which the 4CMenB detergent-extracted OMVs are derived, and the MC58 strain from which the sequence of fHbp contained in the 4CMenB vaccine was derived (Pizza et al., 2000). M01-240364 was selected as a PorB 2 expressing strain to evaluate the impact of different PorB alleles on functional activity. Moreover, the MenB strains M01-240355 (Snape et al., 2013), M07576 (Viviani et al., 2023) and ARG3191 (*Neisseria* isolates database ID: 51591) were selected because they were mismatched for NHBA, NadA, fHbp and PorA P1.7-2,4 variants present in the 4CMenB vaccine. In **Table S2** the genotyping of the 10 MenB strains selected for the bactericidal analysis of antibodies is reported. Our results showed that Gc b-mAbs were highly bactericidal against most MenB strains. Anti-LOS antibodies were able to kill all MenB strains with the exception of NGH38 and ARG3191 (**Fig. 3D, right panel; Table S1 and 2**). Three of these mAbs (01B05, 03N18, 04E05) were more potent than the fourth (02G11) (**Fig.3D, right panel; Table S1**). Anti-PorB b-mAbs also showed broad functionality against MenB strains. Interestingly, while only one anti-LOS exhibited low activity against ARG3191, anti-PorB antibodies were extremely functional against this strain (**Fig. 3D, right panel; Table S1**). Conversely, anti-PorB antibodies showed poor or no activity against MenB strains carrying a PorB class different from the NZ98/254 strain, namely 5/99, NGH38 and M01-240364 (**Fig. 3D, right panel; Table S1**). Antibodies targeting unknown antigens showed the lowest breadth of reactivity against MenB strains and their killing profile was similar to anti-PorB b-mAbs.

**Fig. 3.**
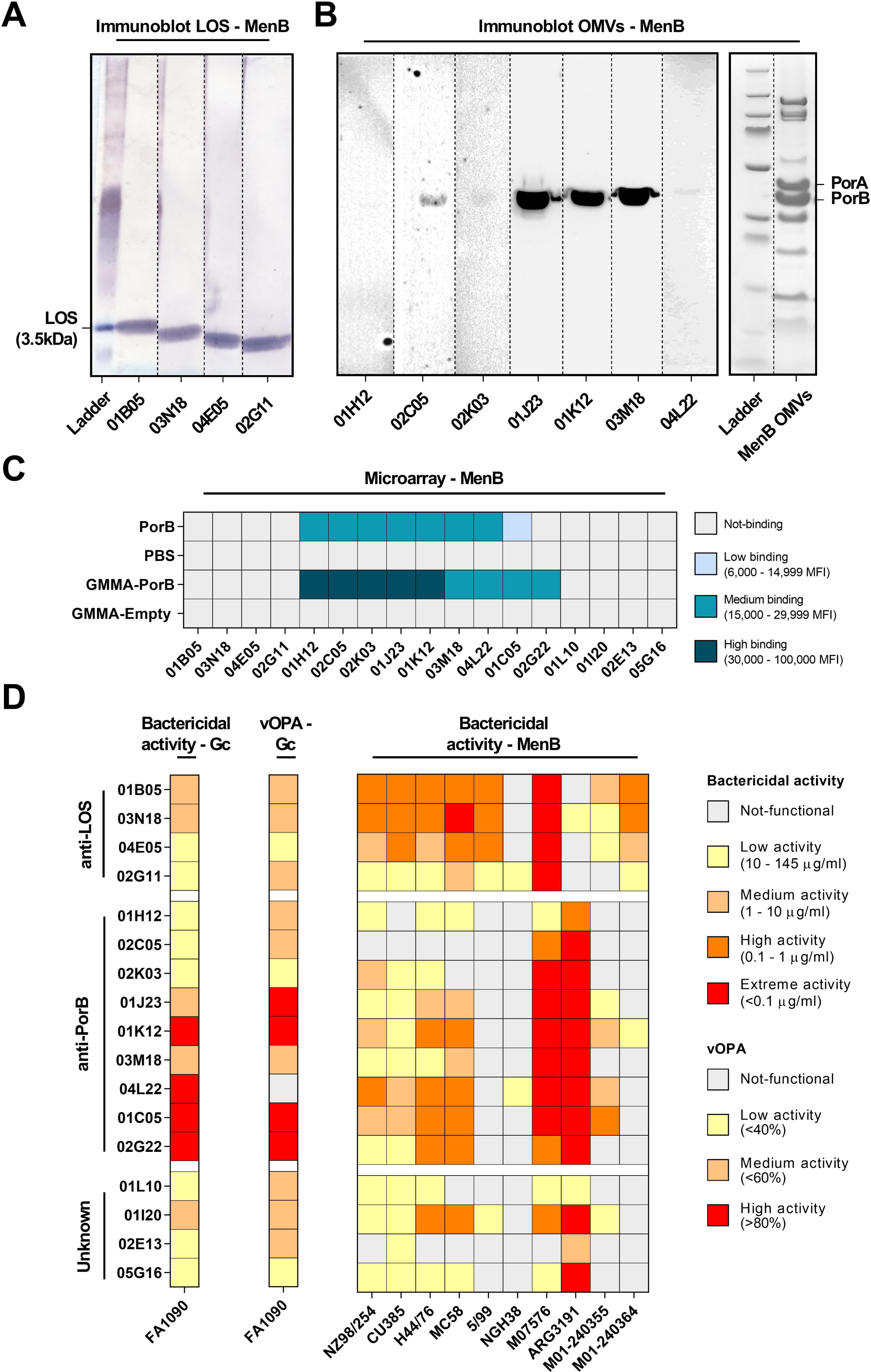
Antigen identification and functional characterization of bactericidal mAbs. (**A-B**) From left to right, immunoblot analysis of anti-LOS (**A**) and anti-PorB mAbs (**B**) on purified meningococcal LOS and on OMVs, respectively. (**C**) The heatmap shows the binding signal intensity of Gc b-mAbs to meningococcal recombinant PorB antigen and to *E. coli* GMMA expressing MenB PorB. (**D**) The heatmap shows on the left the bactericidal activity of b-mAbs against Gc FA1090 strain. The middle panel shows the opsonophagocytic activity of b-mAbs against FA1090. The right panel shows the bactericidal activity of b-mAbs against 10 MenB strains.

### *IGHV4-34* is predominantly used by anti-PorB bactericidal antibodies

We subsequently investigated the genetic characteristics of the b-mAbs. All the genetic features of the 17 b-mAbs are summarized in **Table S3**. Surprisingly, all 9 anti-PorB b-mAbs used the immunoglobulin heavy variable chain (IGHV) 4-34 germline rearranged with the immunoglobulin heavy joining chain (IGHJ) 3-1 (3/9; 33.3%), IGHJ4-1 (3/9; 33.3%) or IGHJ6-1 (3/9; 33.3%). These heavy chains paired with different light chains, most frequently with IGKV3-20 (7/9; 77.8%) (**Fig. S4A**). The b-mAbs targeting unknown antigens also preferentially used the IGHV4-34 germline (3/4; 75.0%) paired exclusively with the IGKV3-20 which accommodated different J genes (**Fig. S4A**). Interestingly, while most of anti-PorB b-mAbs carry the IGHV4-34 paired with IGKV3-20, only two clonal families with two members each were identified (clone ID 6 and 8) (**Table S3**). The remaining anti-PorB antibodies were orphan sequences (i.e. do not belong to clonal families) highlighting the diversity of this class of antibodies. Anti-LOS used mainly the IGHV2-5; IGHJ4-1 (3/4; 75.0%) paired with different light chains (**Fig. S4A**). In addition, 3/4 (75%) anti-LOS b-mAbs derived from the same clonal family (clone ID 1). The low number of anti-LOS b-mAbs and their high clonality means that limited heterogeneity was observed in those recovered in this study. Our analyses revealed that anti-LOS and anti-PorB mAbs use preferentially different heavy and light chain gene rearrangements (**Fig. S4A).** In addition, we evaluated the V gene mutation frequency, heavy chain complementary determining region 3 (HCDR3) amino acidic length, frequency of positively charged and hydrophobic amino acids in the HCDR3. Our analyses showed that anti-LOS b-mAbs had almost 3-fold higher V gene mutation frequencies compared to antibodies targeting PorB or unknown antigens (**Fig. S4B**). Furthermore, anti-PorB and antibodies targeting unknown antigens showed longer HCDR3 (23 – 28 amino acids, compared to 13-15 for anti-LOS b-mAbs), and a higher frequency of positively charged residues compared to anti-LOS b-mAbs (**Fig. S4C-D**). Finally, no major differences in the frequency of hydrophobic amino acids in the HCDR3s were observed among the three groups of antibodies (**Fig. S4E**).

### Immuno-staining of the gonococcal bacterial surface

Following functional and genetic characterization of our panel of 17 Gc b-mAbs, we investigated their binding pattern on the surface of Gc FA1090 (**Fig. 4**). Specifically, three different assays were performed: flow cytometry, to evaluate the percentage of bound bacteria in the whole population, immunofluorescence and immunogold, to profile the binding pattern of each antibody on the surface of single bacteria. Flow cytometry data showed that the majority of anti-LOS antibodies (3/4; 75%) bound 88-92% of FA1090 bacterial population (**Fig. 4A, left panel**). Only the 02G11 antibody showed a lower ability to bind Gc. Antibodies showing strong recognition by flow cytometry also had a very strong signal in immunofluorescence (**Fig. 4A, middle panel**). In addition, immunofluorescence and immunogold analyses revealed that all anti-LOS b-mAbs bind homogenously the entire surface of Gc (**Fig. 4A, middle and right panels; Fig. S5A**). Anti-PorB antibodies showed two different modalities of binding. Flow cytometry data revealed that 6 out of 9 b-mAbs (66.7%) were able to bind over 70% of bacteria while the remaining 3 bound less than 56% of the Gc population (**Fig. 4B, left panel**). A similar pattern was also observed by confocal and electron microscopy (**Fig. 4B, middle panel**). Indeed, almost all antibodies (8/9; 88.9%) bound homogenously the whole surface of the bacterium, while the remaining antibody (01H12) bound discontinuously the surface of Gc, showing higher binding intensity on specific spots of the bacterium (**Fig. 4B, middle and right panels; Fig. S5B**). Antibodies targeting unknown antigens all showed binding to over 67% of the bacterial population by flow cytometry (**Fig. 4C, left panel**) and also showed two different modalities of binding by immunofluorescence and immunogold analyses, similar to what was observed for anti-PorB antibodies. Indeed, two b-mAbs (01I20 and 01L10) bound homogeneously the whole surface of Gc while the remaining two antibodies showed a discontinuous pattern and displayed high-intensity binding only to discrete spots on the Gc surface (**Fig. 4C, middle and right panels; Fig. S5C**).

**Fig. 4.**
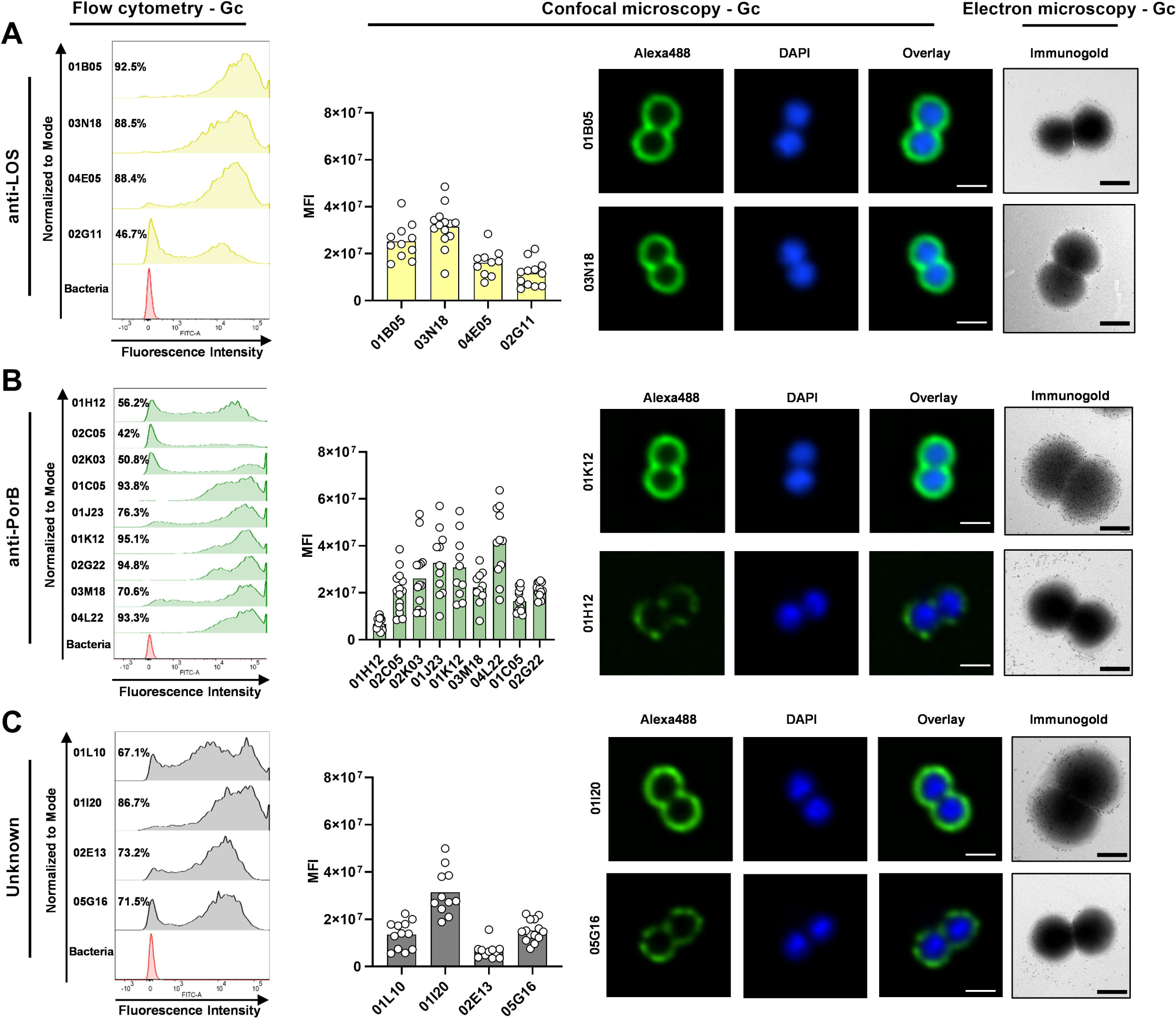
b-mAb binding profiles to Gc FA1090. **(A-C)** Graphs from left to right show flow cytometry histograms, mean fluorescence intensity (MFI) with representative images acquired by confocal microscopy, and images acquired by electron microscopy (immunogold) for anti-LOS (**A**), anti-PorB (**B**) and mAbs targeting unknown antigens (**C**), respectively. For flow cytometry analyses, red, yellow, green, and grey histograms represent negative control, anti-LOS, anti-PorB and mAbs targeting unknown antigens respectively. Immunofluorescence images were acquired with 60X magnification and scale bar reports 1 µm. Representative electron micrographs of immunogold labeling of anti-PorB, anti-LOS and mAbs targeting unknown antigens binding to Gc FA1090, showing their location as indicated by 12 nm gold particles (black dots). Scale bar for immunogold labeling reports 1 µm.

### Epitope characterization of anti-LOS mAbs

Neisserial LOS consists of lipid A anchored into the outer membrane, and an oligosaccharide core composed of two 3-deoxy-D-manno-2-octulosonic acid (KDO) and two heptose residues (Hep1 and Hep2 residues) from which three oligosaccharide chains (α, β, γ) containing glucose (Glc), galactose (Gal), glucosamine (GlcNAc) and galactosamine (GalNAc) can extend (Jennings et al., 1999) (**Fig. 5**). While the γ chain is constitutively expressed in both *Neisseria* species, glycan extensions of the α-chain from Hep1 and the β-chain from Hep2 are controlled by a series of phase variable LOS glycosyltransferase (*lgt*) genes (Jennings et al., 1999) resulting in variable length of sugar chains between and within strains. Phase variable *lgtA, lgtC* and *lgtD* encode glycosyl transferases involved in the elongation of the LOS α-chain, while *lgtG* mediates LOS β-chain extension. LOS variation in MenB leads to 12 immunotypes (L1-L12) (Mubaiwa et al., 2017) all carrying an α-chain of variable length (from 2 to 4 sugars) and only two of them (L2, L5) bearing a β-chain with one glucose. The NZ98/254 MenB strain from which 4CMenB OMV component is prepared belongs to L1 and L3,7,9 immunotypes (Findlow et al., 2007). Although there is no LOS typing scheme defined for Gc, all strains contain an α-chain that can be composed of 2, 3, 4 or 5 sugars, while the β-chain, composed of a lactose, can be present or absent. To shed light on the specificity of the anti-LOS b-mAbs discovered here, we utilized a library of 8 LOS mutants derived from the MS11 4/3/1 strain, in which the 4 variable *lgt* genes were genetically fixed “ON” or “OFF” (Chakraborti et al., 2016). Each of these strains mainly expresses one of the Gc LOS structures, which facilitated the identification of epitopes recognized by the b-mAbs binding to the bacterial surface (**Fig. 5; Fig. S6A**) (Chakraborti et al., 2016). Specifically, the MS11 mutant strains presented 8 different LOS structures, 4 carrying both α- and β-chains (2HexG+, 3HexG+, 4HexG+ and 5HexG+) and 4 carrying only the α-chain (2HexG-, 3HexG-, 4HexG- and 5HexG-), where G+ and G-refer to mutants in which the status of *lgtG* is fixed “ON” or “OFF” respectively, while the designation 2Hex, 3Hex, 4Hex and 5Hex refers to mutants expressing 2, 3, 4 and 5 sugars respectively in the α-chain. LOS structures 2HexG-, 3HexG- and 4HexG-are known to be shared between Gc and MenB (Mubaiwa et al., 2017). Immunoblotting analysis on the LOS MS11 mutants revealed that all anti-LOS b-mAbs bound to the 2HexG-, 3HexG-, 4HexG- and 5HexG-structures, while the presence of the β-chain impaired bacterial recognition (**Fig. 5; Fig. S6A**). The data indicate that the minimal structure required for these antibodies to bind the LOS was the 2HexG-, characterized by a single α-chain composed of two hexoses and by the absence of the β-chain. To validate the obtained immunoblot data, the 4 anti-LOS b-mAbs were tested for binding on a microarray containing the 4CMenB OMV present in the vaccine, and additional OMVs from 13 MenB strains known to carry different LOS structures (**Fig. S6B; Table S2**) (McLeod Griffiss et al., 2000; O’Connor et al., 2008). Microarray data showed that anti-LOS b-mAbs were able to bind 83.3% of MenB strains with absent or phase OFF *lgtG* gene, while they were not able to recognize the MenB strains with *lgtG* gene ON (**Fig. S6B; Table S2**).

**Fig. 5.**
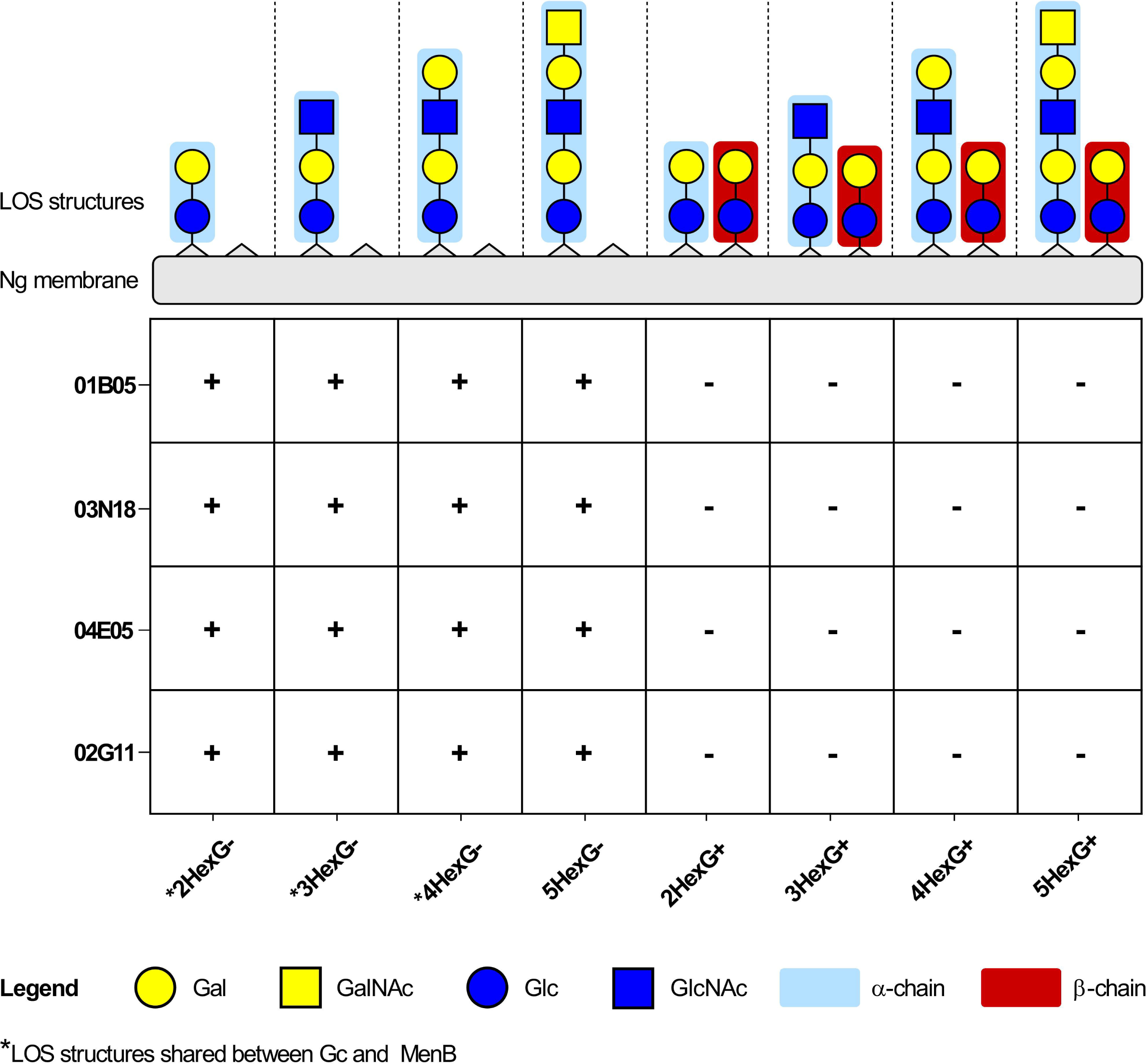
Characterization of the anti-LOS mAbs on Gc MS11 LOS mutants. The graph shows the positive (+) and negative (-) binding of the 17 Gc b-mAbs to the purified LOS structures of 8 MS11 mutants.

### Epitope characterization of anti-PorB b-mAbs

PorB is a voltage-gated pore found in the outer membrane as a homotrimeric B-barrel. Each PorB monomer of 32 to 35 kDa is composed of 16 transmembrane-spanning regions and 8 extracellular loops (Chen and Seifert, 2013). The MenB (NZ98/254) and Gc (FA1090) PorB regions recognized by 01K12, the anti-PorB b-mAb showing the highest functional activity against Gc, were investigated by Hydrogen/Deuterium exchange Mass Spectrometry (HDX-MS) and by *in silico* Molecular Docking. 4CmenB and Gc detergent-extracted OMVs were labeled with deuterium in the presence or absence of 01K12 and the level of H/D exchange was monitored on 107 and 78 PorB peptides respectively covering more than 98% of the MenB and Gc protein sequences. Upon binding of the b-mAb to 4CmenB OMVs, reduction in deuterium incorporation was observed on 21 overlapping peptides, which defined an epitope involving loop 5 (V186-H194), loop 6 (D215-L235) and loop 7 (S255-L261) (**Fig. 6A; Fig. S7**). Interestingly, Gc PorB showed a different mode of binding, with reduction of deuterium incorporation in loop 1 (T21-S26), loop 3 (N94-A106 and I119-Y132), loop 4 (S143-Y161), loop 5 (F180-T196) and loops 6-7 (Y235-D284) (**Fig. 6B**). *In silico* docking analysis was performed considering the 01K12 b-mAb and MenB PorB, and Gc PorB respectively. In particular, for antibody structural prediction the DeepAb, an artificial intelligence (AI) algorithm specific for antibody modeling that provides a highly confident estimation of the CDR3 region was used, while the paratope region was predicted with Paragraph (Chinery et al., 2023) on the generated models. Additionally, AlphaFold2 (Jumper et al., 2021) was adopted to predict the NZ98/254 MenB PorB and FA1090 Gc PorB structures (**Fig. 6C-D**). By comparing MenB PorB and Gc PorB 3D models a significant difference was observed in the length and structural features of Loop 5, that appeared short and disordered in MenB PorB and instead 13 residues longer and well-structured in Gc PorB (**Fig. 6E**). In Gc PorB, loop 5 belonging to one monomer was oriented towards the adjacent monomer influencing the conformation of the overall epitope region, while this was not observed for MenB PorB. On the basis of these observations, docking epitope definition was confined to the monomer for MenB PorB and to the trimer of Gc PorB. Conformational structural analysis on 5 models obtained for Gc PorB mainly revealed 5 different orientations for loop 5 with a Root Mean Square Deviation (RMSD) of 8 Å (**Fig. S8A**; orientation 1-5 in **Table S4**) while for 5 MenB PorB models only three 3 possible orientations for loops 5-8 with an RMSD of 4 Å (**Fig. S8B**; orientation 1-3 in **Table S5**). These ensembles of PorB orientations were used as a starting point for docking analysis. The docking results showed good agreement with HDX-MS data. Indeed, binding analysis on the lowest energy docking pose for MenB PorB/01K12 (-42,57 Haddock Unit, table S2) and Gc PorB/01K12 (-49,324 Haddock Unit-Table S3) revealed the same loop binding pattern observed experimentally (**Fig. 6**).

**Fig 6.**
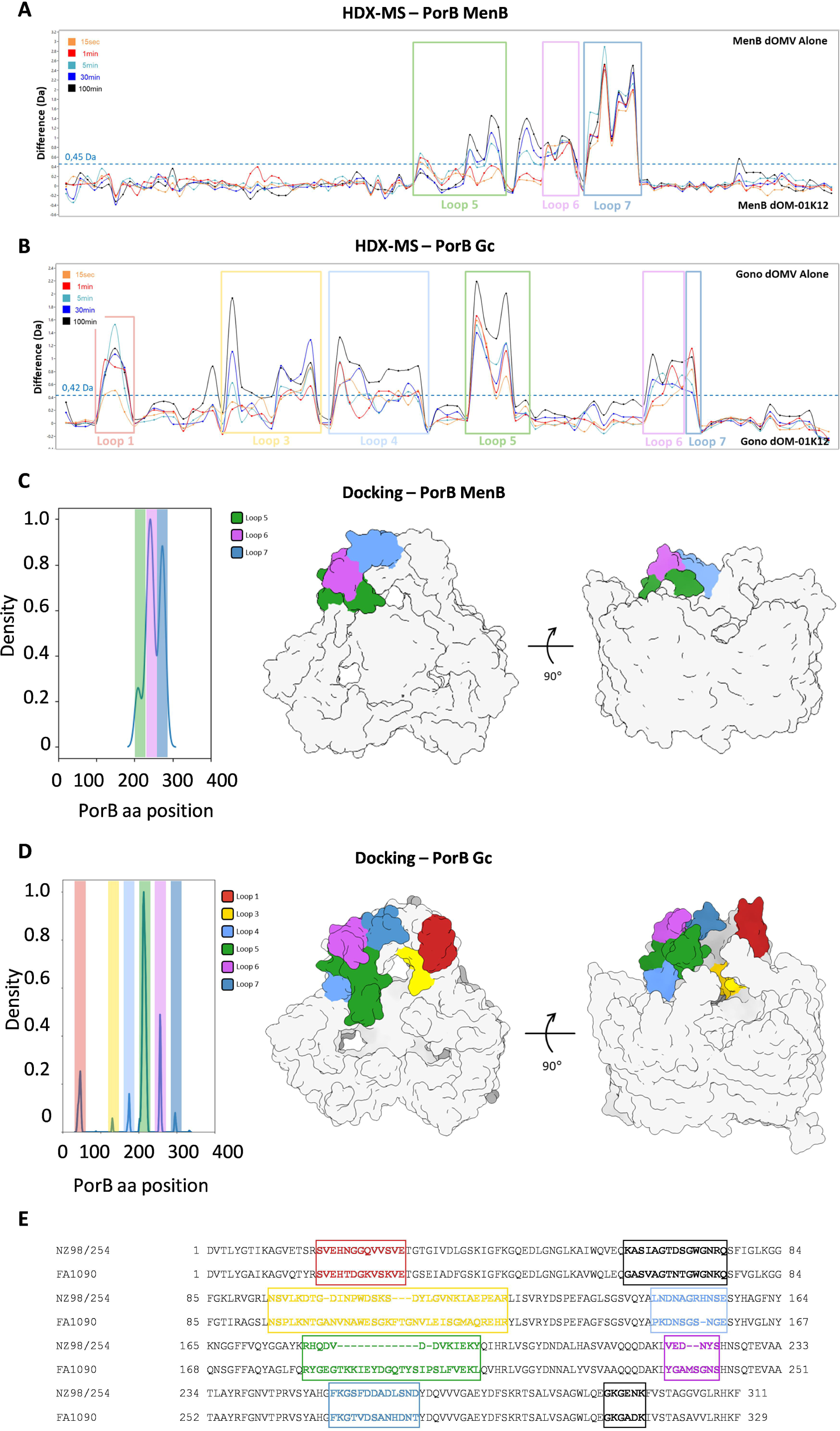
Epitope characterization on trimeric MenB and Gc PorB. (**A-B**) MS epitope mapping of b-mAb 01K12 on MenB PorB (**A**) and Gc PorB (**B**). Differences in deuterium uptake (Y-axis) of PorB in absence and presence of b-mAb 01K12 for the 107 (**A**) and 78 (**B**) identified peptides during time course from 15 sec to 100 min. Individual peptides are arranged along the X-axis starting from the N-terminus to the C-terminus of the protein. Positive value in Y-axis indicate protection in deuterium incorporation in presence of b-mAb. Dotted lines indicate the 98% CI. (**C-D**) Molecular docking results of b-mAb 01K12 on MenB PorB (**C**) and Gc PorB (**D**). The Antigen/Antibody contact probability distribution was calculated on the lowest PorB/01K12 energy cluster. On the Y-axis is reported the contact density distribution and on the X-axis the residues number relative to PorB. Plots are colored according to the loops in contact with PorB: loop 1 (red), 3 (yellow), 4 (light blue), 5 (green), 6 (purple), and 7 (dark blue). Graphical representation of the relative loop position with respect to the β-barrel region. (**E**) Sequence alignment of MenB (NZ98/254) and Gc (FA1090) PorB. PorB loops are identified by bold residues. Colored loops were identified as relevant in MenB and Gc PorB HDX-MS epitope mapping.

### Protection in a low genital tract mouse model of infection

An *in vivo* lower genital tract (LGT) mouse model of infection of female BALB/c mice was applied to assess the protective efficacy of 01K12 against the Gc strain FA1090 (**Fig. 7A**). 01K12 was selected as it showed the highest bactericidal potency against Gc and broad recognition of clinical Gc strains and MenB strains, suggesting resistance to multiple mutations on the PorB antigen (**Fig. 3D and Fig. S3**). To allow colonization of the mice by the challenge gonococcal strain mice were treated with estradiol and antibiotic prior to infection. The efficacy of mAb 01K12 was assessed by daily intravaginal administration throughout the course of the experiment. For this study, 42 mice were divided into three groups (14 animals each). The first group was composed by animals treated with 0.5 µg/day 01K12 for one week. The second and third groups were the Placebo and HexaBody isotype control groups, which received a saline solution and an anti-respiratory syncytial virus (RSV) antibody at the concentration of 0.5 µg/day for one week (**Fig. 7A**). Animals were challenged with 2×10^6^ colony forming units (CFU) of Gc delivered intravaginally 2 hours post-antibody and saline solution administration. Subsequently, vaginal swabs and CFU counts were performed daily throughout the seven days of the study. The percentage of infected mice and bacterial burden, measured as area under the curve of Gc CFU over time, were used to evaluate the efficacy of the mAb. Mice treated with 01K12 exhibited a significantly faster clearance rate compared to the saline (p=0.0235) and isotype (p=0.0222) groups. In addition, our results showed that only 36% of mice receiving 01K12 were colonized on day 7 compared to 71% and 78% of mice receiving saline or HexaBody isotype control, respectively (**Fig. 7B**). Finally, significantly decreased bacterial burdens in the group receiving 01K12 were observed compared to the saline and HexaBody isotype control groups (**Fig. 7C**).

**Fig. 7.**
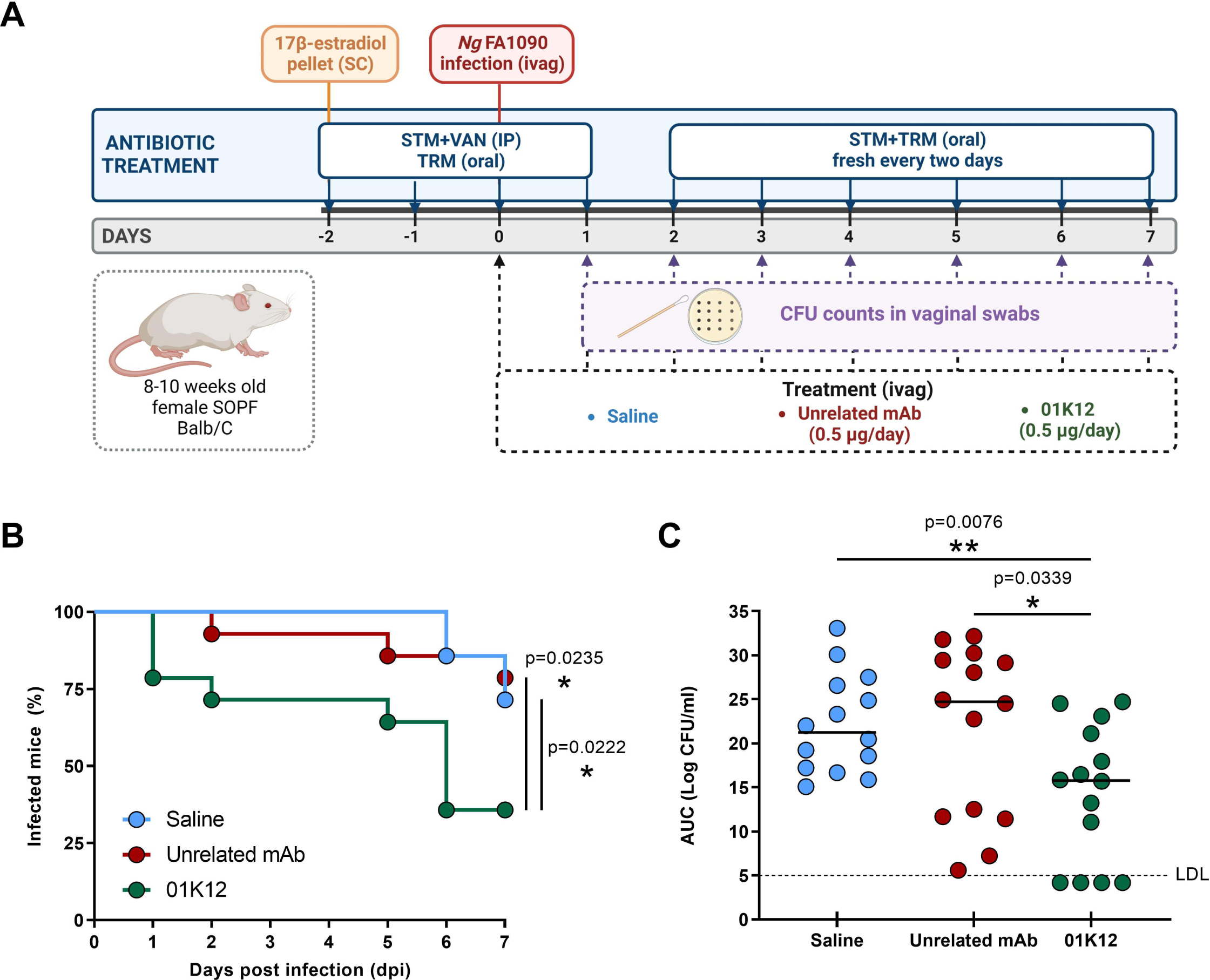
Prophylactic efficacy of 01K12 in a mouse LGT model of Gc infection. (**A**) Schematic representation and timelines of therapeutic studies performed in SOPF-BALB/c mice to assess the protective efficacy of 01K12. Mice were initially treated with streptomycin (ST), vancomycin (VAN) and trimethoprim (TRM) and then challenged intravaginally (ivag) with Gc FA1090 on day 0 (D0). Average bacterial burden was measured daily in vaginal secretions after infectious challenge on D1. (**B**) Time to bacterial clearance is shown using Kaplan-Meier curves, which display the percentage of each group with detectable vaginal CFU, measured daily after infectious challenge on D1. (**C**) Overall bacterial titer in each animal was assessed using the area under the curve (AUC) analysis, calculated from the distribution of the CFUs over time. AUC values for each group were compared using the Mann Whitney U test. Significances are shown as Up < 0.05 and UUp < 0.01.

## DISCUSSION

Gc is one of the bacterial pathogens which is resistant to most antibiotics and for which no alternative medical tools such as vaccines or monoclonal antibodies are currently available. In this work, we were prompted by recent reports showing that the licensed 4CmenB vaccine induced protection against Gc to investigate the nature of the immune responses resulting from 4CmenB vaccination and to clone and identify MBCs from vaccinees which produced antibodies able to kill Gc. Surprisingly, the antibodies identified recognized only PorB protein and LOS among the many antigens shared by Gc and MenB. These antigens were already known to elicit bactericidal activity but are not considered as priority antigens in MenB or Gc vaccines. The OMV component used in the licensed MenB vaccine had been included in the vaccine largely because they carry the immunodominant PorA antigen which, despite being antigenically variable, induces strong bactericidal response against homologous strains of MenB, but is not present in Gc (Humphries et al., 2006; Sacchi et al., 1998; Tondella et al., 2000). PorB, instead, is the most abundant antigen on the gonococcal surface, it is densely present on the bacterial surface and is readily accessible as a target of adaptive immunity. However, PorB has been studied more for its immune evasion properties than as a vaccine antigen (Jones et al., 2023). Indeed, PorB is known to bind C4BP and factor H and to contribute to the serum resistance of Gc (Ram et al., 2001) and has other immunomodulatory properties. Monoclonal antibodies obtained by immunizing mice with gonococcal strains had been shown to be bactericidal and opsonic in the presence of complement and able to protect epithelial cells against Gc invasion (Heckels et al., 1987; Joiner et al., 1985), however the 70% amino acid sequence similarity between PorB of MenB and Gc suggested a limited cross-protective role against gonococcus. Surprisingly, we demonstrated that MenB PorB was able to elicit cross-functional antibodies against Gc. Indeed, this is the first study reporting the isolation from humans of Gc bactericidal anti-PorB antibodies from subjects immunized with MenB OMVs. More importantly, the most potent PorB antibody has been demonstrated to be sufficient for protection against urovaginal challenge of gonococcus in the mouse model. This has been shown before for the 2C7 LOS antibody, isolated after immunization of mice (Gulati et al., 2019a). The anti PorB antibodies identified show a great variability in PorB recognition as assessed by immunoblot and immunofluorescence. This can probably be explained by the fact that PorB requires RmpM and other proteins to be stabilized at the outer membrane (Jansen et al., 2000; Marzoa et al., 2010) and that several of the epitopes identified may recognize conformations stabilized by these interactions. Another important vaccine target is represented by LOS. On a molar basis, LOS is the most abundant gonococcal outer membrane molecule and plays a key role in many facets of pathogenesis (Ram et al., 2018). Regarding the similarity or LOS between the two *Neisseria* species, it is known that MenB and Gc share common LOS structures that could elicit cross-reactive antibodies. Shared LOS structures carry an α-chain of variable length, while lack the β-chain (Mubaiwa et al., 2017; Ram et al., 2018). In particular, the NZ98/254 strain predominantly expresses the L1 and L3,7,9 structure shared with Gc bearing an α-chain with 3 and 4 sugars respectively. We found that anti-LOS b-mAbs elicited by MenB OMVs were able to recognize Gc strains carrying LOS structures with α-chain of variable lengths (2 to 5 sugars), likely through binding to the 2Hex-structure as common minimal epitope therein. Importantly the LOS b-mAbs can recognize the Lacto-N-neotetraose (LNnT) epitope (also referred as L3,7,9) (Tsai and Civin, 1991) which is essential for LOS-mediated adherence and invasion and, upon sialylation, promotes Gc serum resistance. Those LOS structures are complementary to the ones recognized by the 2C7 mAb, the anti-gonococcal immunotherapeutic previously identified by the Sanjay Ram group, which binds structures containing a β-chain (Yamasaki et al., 1999). The fact that antibodies to LOS (2C7) and PorB (01K12) are protective against gonococcal infection in the urovaginal niche, confirms that humoral responses may be sufficient for protection against gonorrhea. It is also interesting that both of these antigens are key molecular interactors in immune evasion against complement, binding complement down-regulators such as factor H and C4BP suggesting that counteracting mechanisms of immune evasion may be essential in immune interventions against gonococcus. The identification of LOS and PorB as antigens inducing cross-functional antibodies paves the way for the development of vaccines and therapeutics against Gc and sheds light on the broad protection conferred by OMVs contained in the 4CMenB vaccine against Gc and MenB which extends beyond what had been predicted with known antigens (Marjuki et al., 2019; Martinón-Torres et al., 2021). Finally, our study shows that isolating human mAbs from vaccinated or infected people, a practice which has become very popular for viral infections during the Covid-19 pandemic, can be a useful strategy also to identify key protective antigens among the hundreds of potential targets present in bacteria. The rapid identification of the key protective antigens can be very useful to develop vaccines and therapeutics to address antimicrobial resistance, which is one of the most important medical challenges of our times.

### Limitations of the study

A limitation of this study is that *in vitro* neutralization and *in vivo* protection in the gonococcus LGTI mouse model of infection cannot be fully predictive of the behavior of the same antibody in humans and therefore the real benefit of described antibodies can only be assessed in clinical studies.

## Acknowledgments

This work was funded by the European Research Council (ERC) advanced grant agreement no. 787552 (vAMRes). The authors thank Alfredo Pezzicoli for supporting confocal microscopy acquisition, Erica Borgogni and Elisa Faenzi for binding analysis, Giacomo Vezzani and Chiara Nocciolini for b-mAb expression, Sara Marchi for b-mAb purification, Silvia Guidotti for sequencing, Silvia Principato for contributing to MenB immunotyping, Marco Tortoli, Stefania Torricelli and Paolo Perfetti for assistance during *in vivo* studies, Nicola Pacchiani for supporting the analysis of antibody repertoire, Alessandro Muzzi and Alessandro Brozzi for genome assembly, Alessia Biolchi and Michela Brazzoli for manuscript revision. For kindly providing meningococcal serogroup B strains, the authors are grateful to: Xin Wang and Henju Marjuki and Jarad Schiffer (CDC, Centers for Disease Control and Prevention, Atlanta, GA, USA); Richard Moxon (University of Oxford, Oxford, United Kingdom); Diana R. Martin (Institute of Environmental Science and Research, Porirua, New Zealand); Ray Borrow (Health Security Agency, Manchester, United Kingdom); Dominique A. Caugant (NIPH, Norwegian Institute of Public Health, Oslo, Norway) and Adriana Efron (Instituto Nacional de Enfermedades Infecciosas-ANLIS “Dr. Carlos G. Malbrán”, Buenos Aires, Argentina) on behalf of the Argentinian National Laboratories Network (NLR). The BG (Bristol group) strains were kindly provided by Dr. Darryl Hill at the University of Bristol, United Kingdom.

## Author contributions

O.F., E.A. and R.R. conceived the study. P.F., C.C., S.G. and R.N. enrolled 4CMenB vaccinees in the study. E.A., I.P., M.T., S. Tavarini. and C. Sammicheli performed PBMC isolation and single cell sorting. S. Stazzoni, M.T., V.A. and G.A. performed B cell binding and bactericidal screening. S. Stazzoni., M.T., V.A., G.A., E.M., P.M., M.D.V., G.T. and C.L. retrieved antibody sequences, expressed and purified recombinant antibodies. E.A. and A.R. performed B cell repertoire analyses. F.C. performed immunoblot on MenB and Gc LOS. V.V. performed microarray analyses. S. Tomei and L.S. performed classical SBA on GC and MenB strains. L.E.F. and S.F. performed HDX-MS experiments. F.V., D.C. and C. Sala produced and analyzed visual opsonophagocytosis data. V.A., E.M. and P.M. performed immunoblot on MenB dOMVs. F.F. and C.T.B. performed immunofluorescence assays. F.G. performed immunogold staining. M. Bodini. analyzed MenB strains. M.A. performed docking analysis on MenB and Gc PorB. A.S. performed *in vivo* experiments. S. Stazzoni, M.F., M.T., V.V., F.C., L.E.F., M.A., L.S., B.B., I.M., I.D., O.F., E.A. and R.R. wrote the manuscript. All authors undertook the final revision of the manuscript. M.F., M. Bardelli, I.M., I.D., O.F., E.A. and R.R. coordinated the project.

## SUPPLEMENTARY FIGURE LEGENDS

**Fig. S1.**
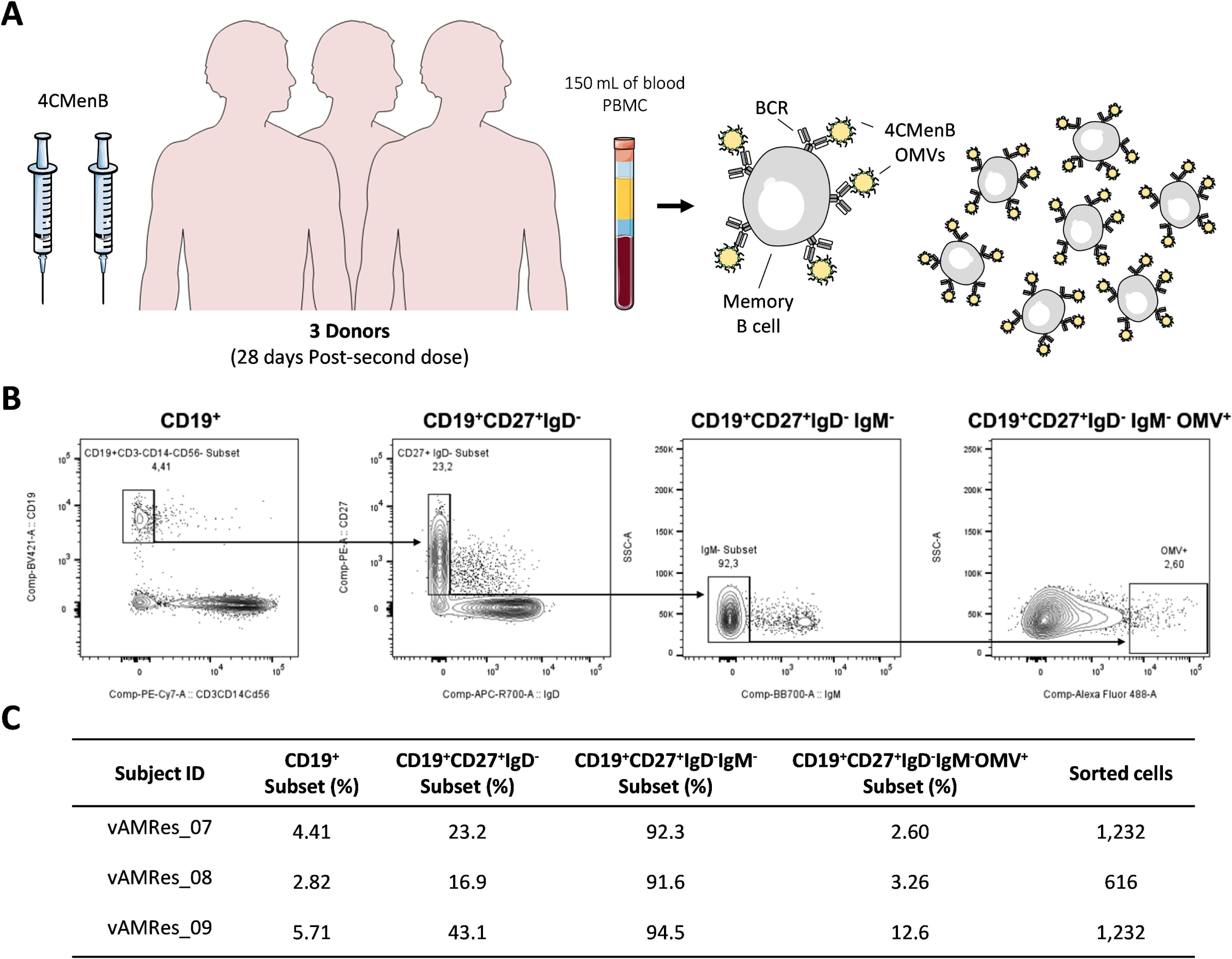
Gating strategy used for single cell sorting of MBCs. (**A**) Schematic representation of cohort and strategy to isolate 4CMenB OMV^+^ MBCs. (**B**) Starting from top left to the right panel, the gating strategy shows: CD19^+^ B cells; CD19^+^CD27^+^IgD^-^; CD19^+^CD27^+^IgD^-^IgM^-^; CD19^+^CD27^+^IgD^-^IgM^-^OMV^+^ B cells. (**C**) Table shows the overall B cell subset frequencies and number of sorted cells.

**Fig. S2.**
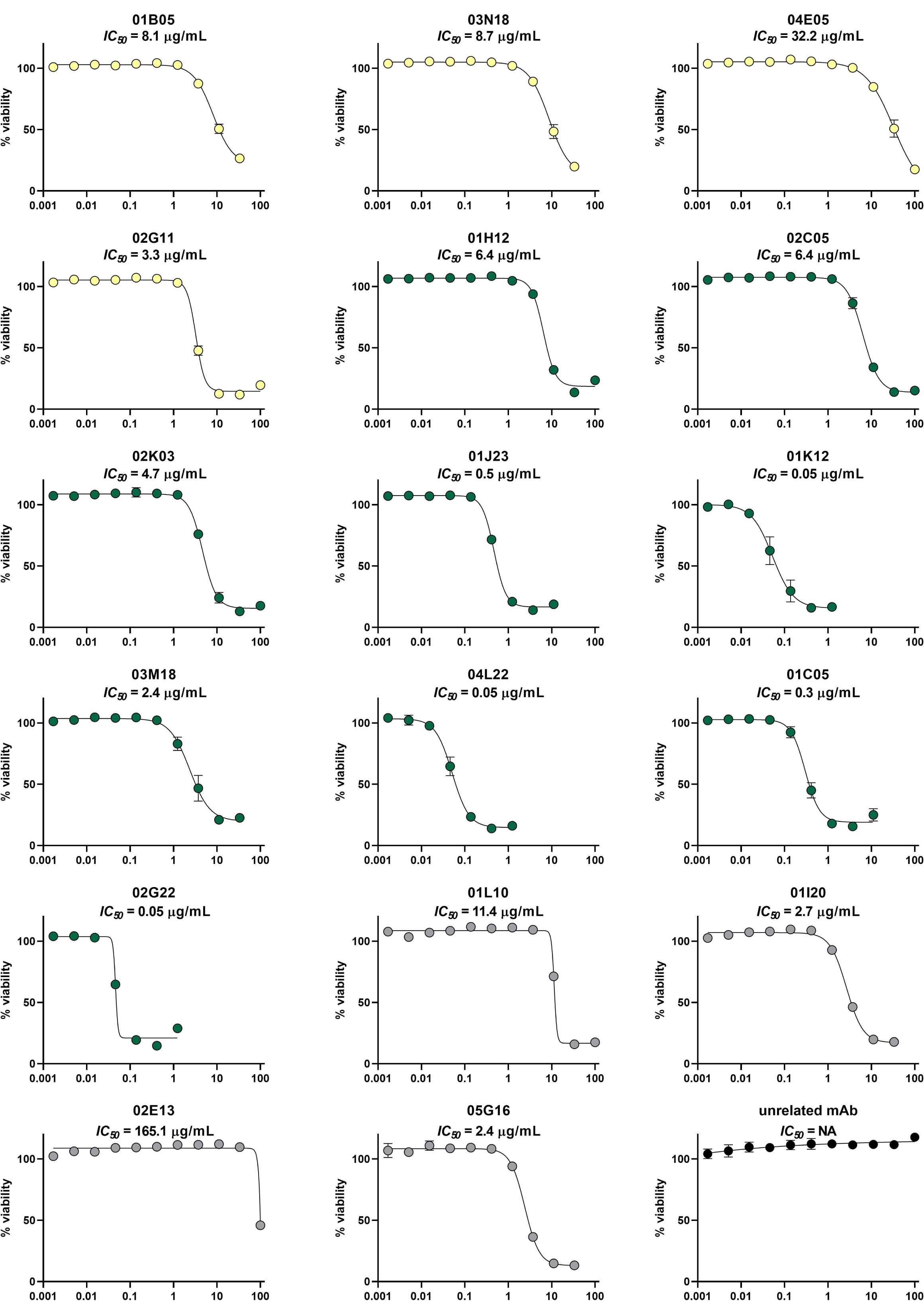
Bactericidal activity of HexaBody mAbs by R-ABA against Gc FA1090 strain. mAbs are sorted by color. Anti-LOS mAbs are displayed as yellow dots, anti-PorB mAbs as green dots, mAbs recognizing unknown antigens as grey dots and an unrelated mAb (negative control) as black dots. Potency evaluation (IC_50_) was assessed by R-ABA against FA1090 (ATCC) strain. Error bars display the standard deviation over two replicates.

**Figure S3.**
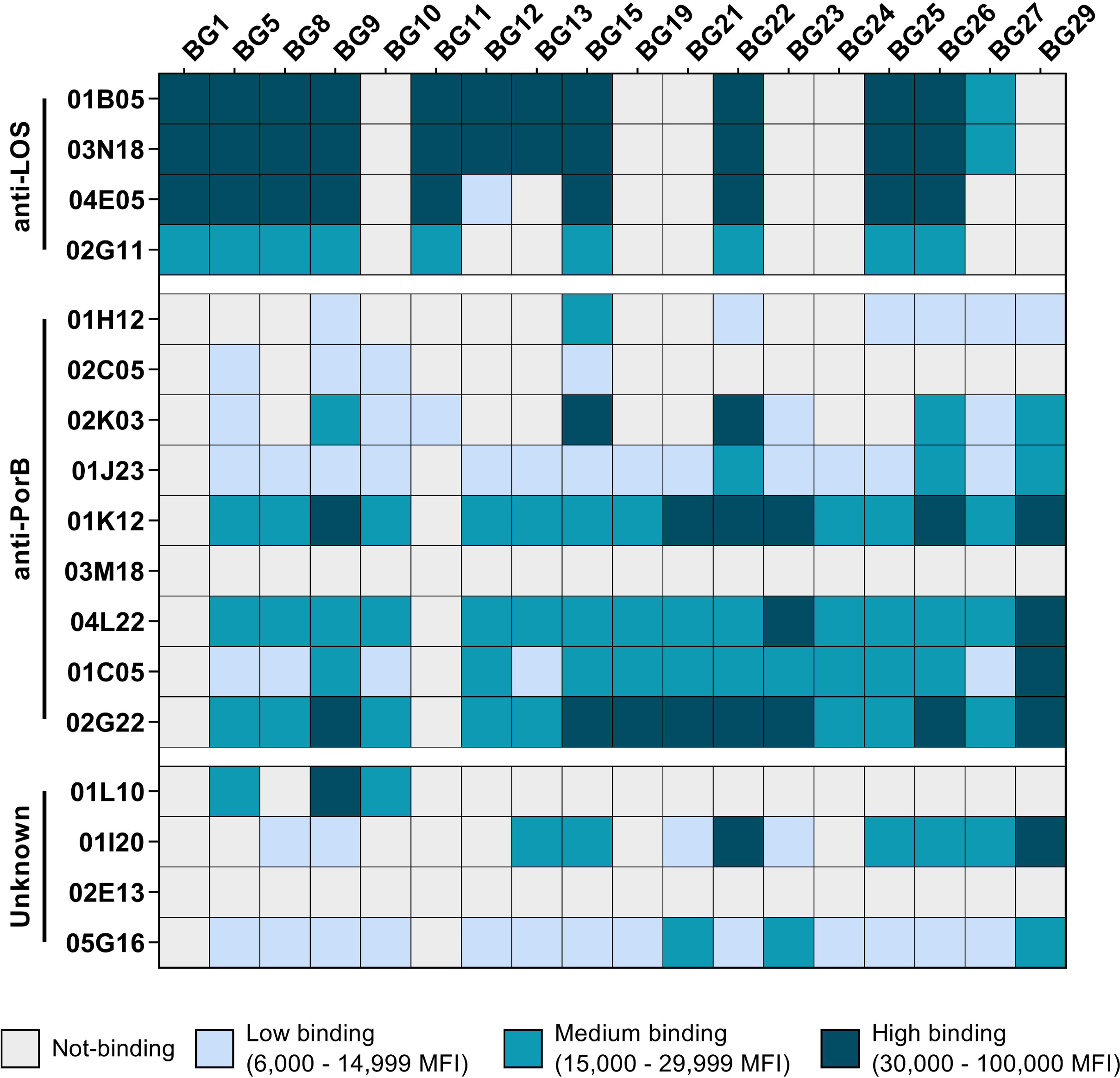
Binding reactivity of b-mAbs to OMVs derived from different Gc clinical isolates. The heatmap shows the binding reactivity of b-mAbs to OMVs derived from 17 low-passaged Gc clinical strains.

**Fig. S4.**
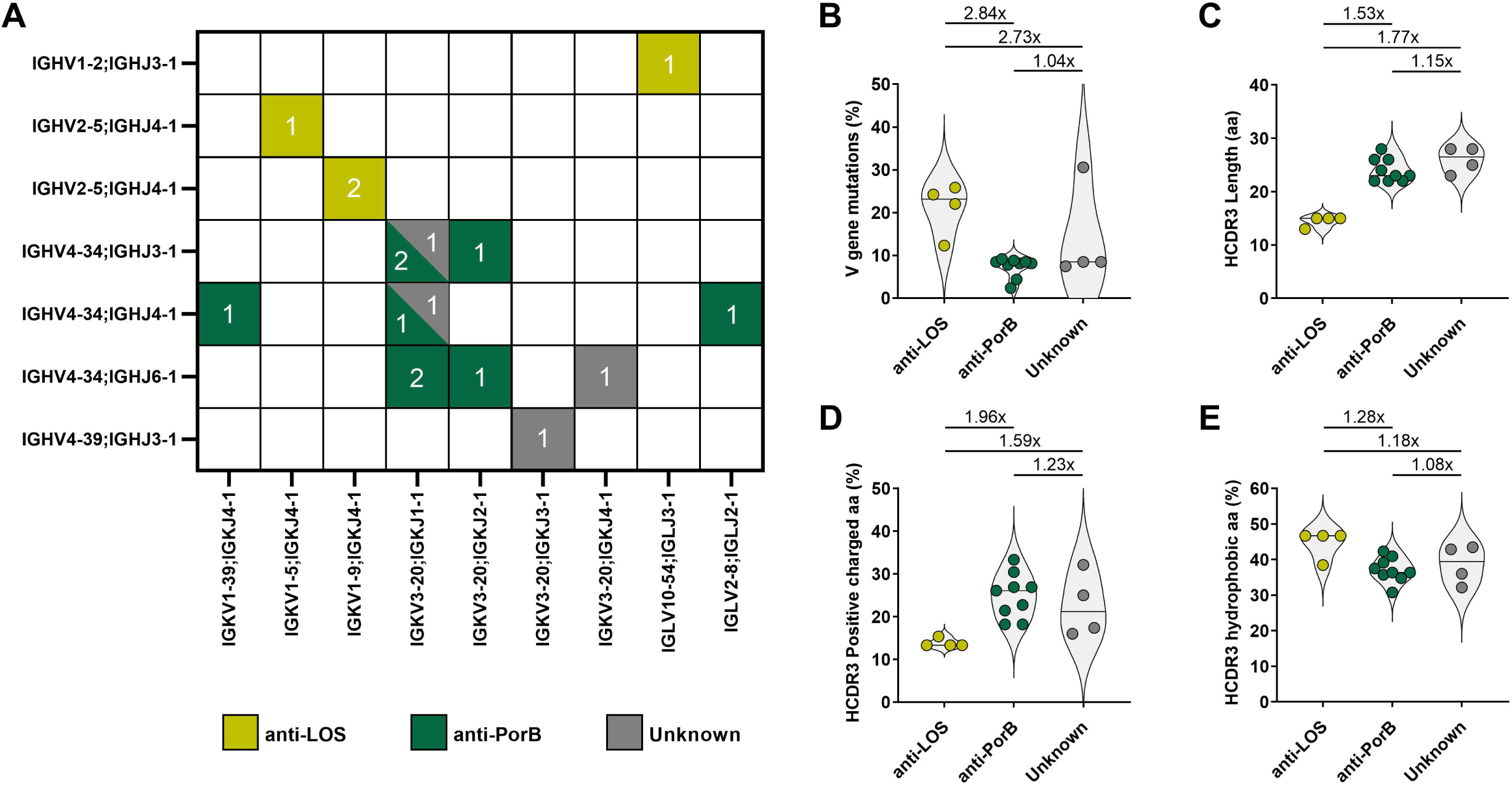
Genetic analysis of bactericidal mAbs. (**A**) Numbers indicate IGHV and IGK/LV matches among mAbs. The two most common combinations are IGHV4-34 pairing with IGKV3-20 for anti-PorB and unknown mAbs, while anti-LOS mAbs showed preferential IGHV2-5 pairing with IGKV2-5. (**B**) V gene mutation is indicated by violin plots as percentage of mutation with respect to the reference germline sequences. (**C-E**) Violin plots indicate the HCDR3 length (aa) (**C**), HCDR3 positive charged aa (**D**) and HCDR3 hydrophobic aa (**E**).

**Fig. S5.**
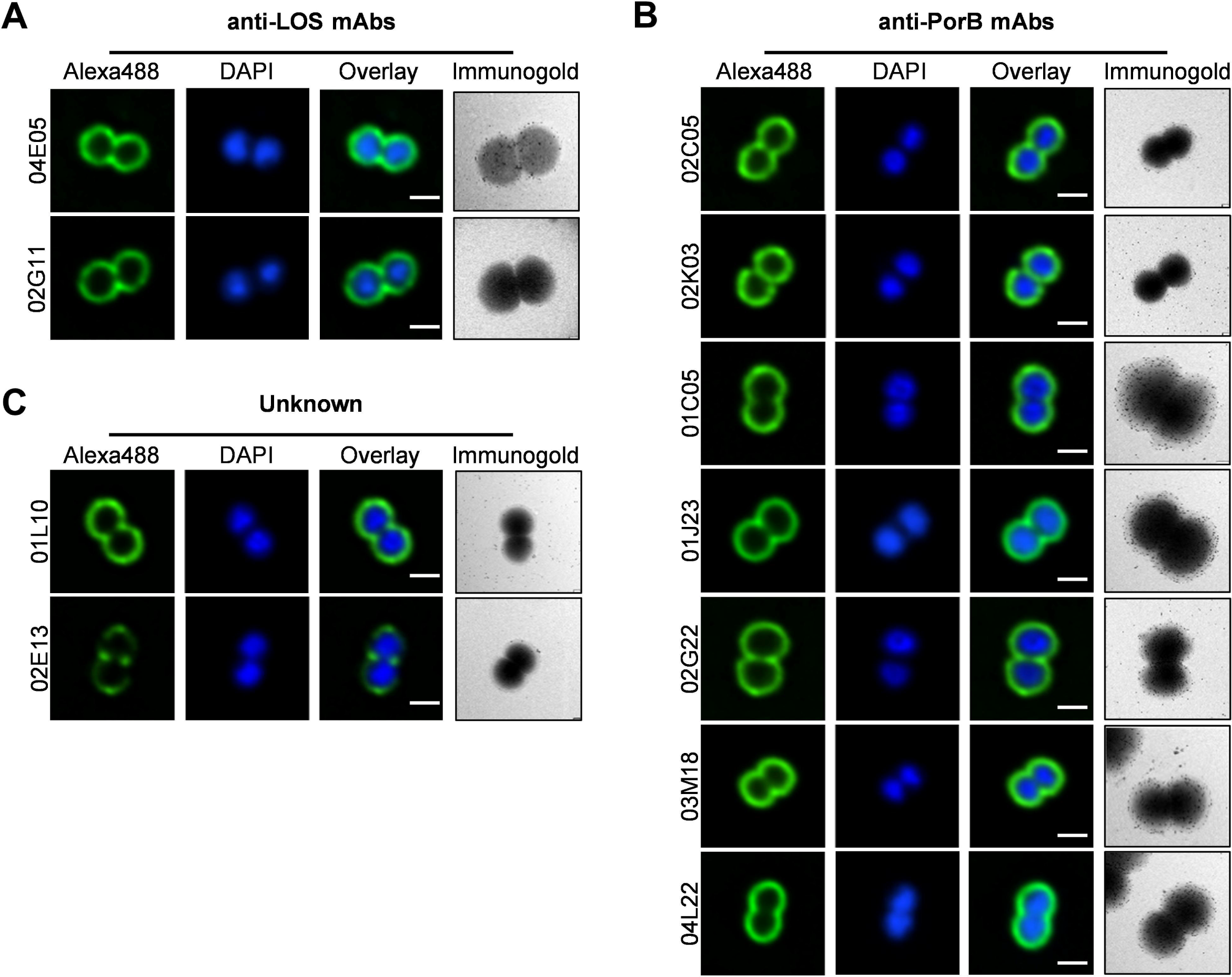
Immunoprofiling of mAbs. (**A-C**) Immunofluorescence and immunogold images obtained for anti-LOS (**A**), anti-PorB (**B**) and unknown (**C**) mAbs. Confocal fluorescent images were acquired with 60X magnification and scale bar reports 1 µm. Immunogold labeling of mAbs showing their location as indicated by 12 nm gold particles (black dots). Scale bar for immunogold labeling reports 1 µm.

**Fig. S6.**
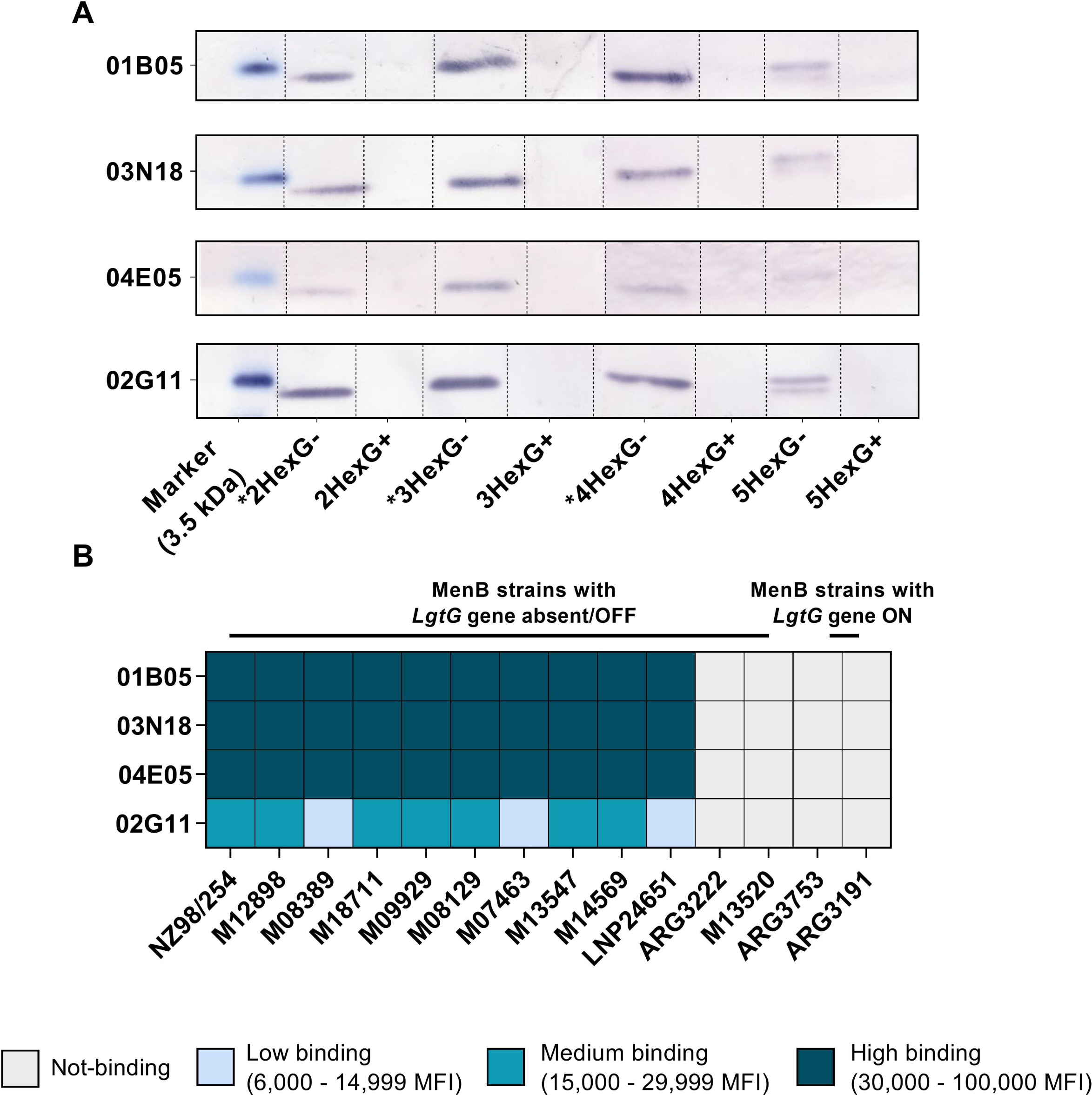
Binding profile of anti-LOS b-mAbs to LOS of Gc MS11 mutant strains and to OMVs derived from a panel of MenB strains. (**A**) Immunoblots showing the binding of the 4 Gc b-mAbs to the purified LOS structures of 8 MS11 mutants. (**B**) The heatmap shows the binding of 4 b-mAbs to 14 MenB strains. Gray, light blue, aquamarine and dark aquamarine identify non-binding, low-, medium- and high-binding, mAbs respectively.

**Fig. S7.**
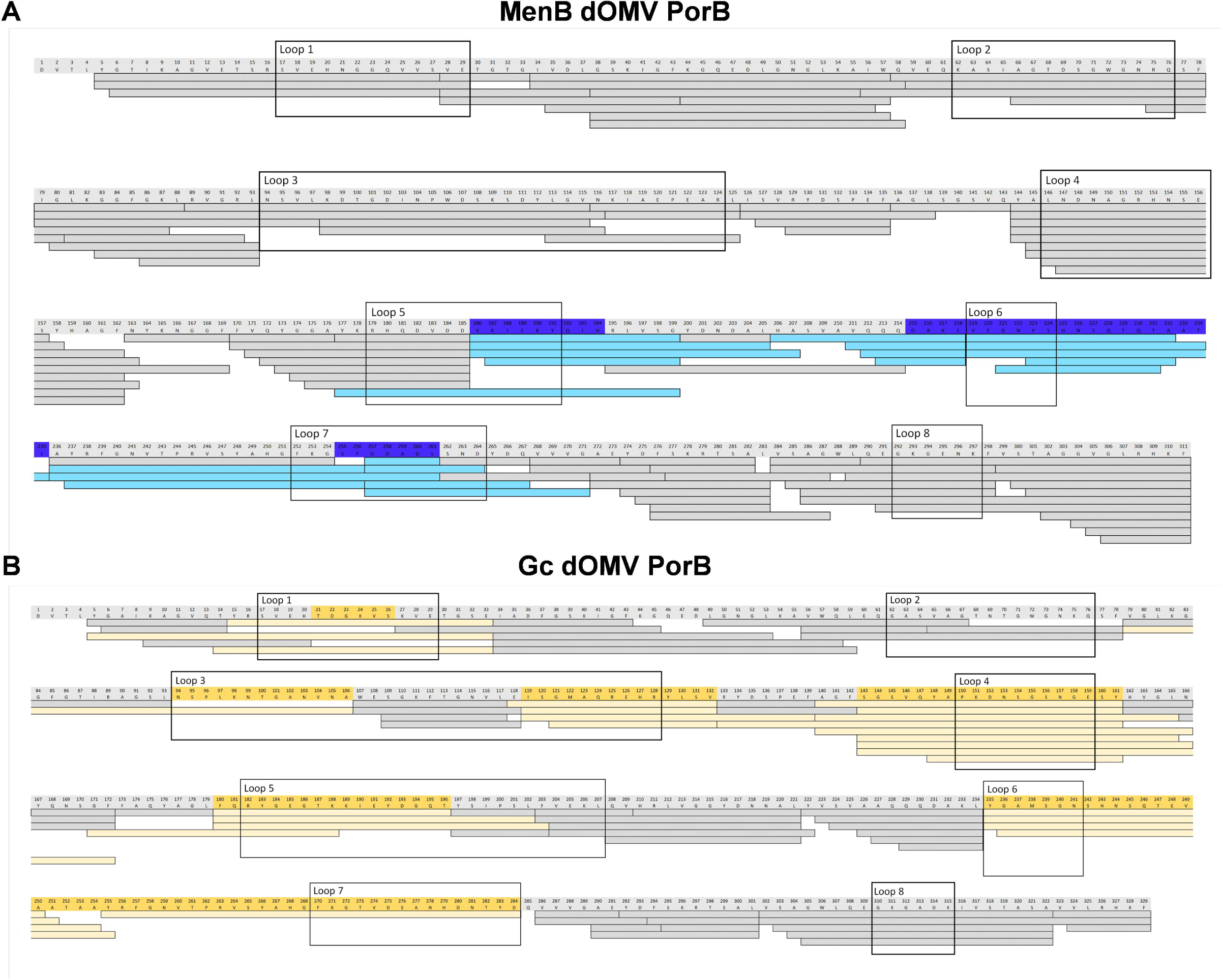
MenB and Gc dOMV-Embedded PorB epitope recognized by b-mAb-01K12. (**A-B**) HDX-MS epitope mapping of b-mAb 01K12 on MenB (**A**) and Gc (**B**) dOMV-embedded PorB The H-D exchange was monitored on 107 (**A**) and 78 (**B**) peptides, covering more than 98% of the protein sequence. No significant differences in deuterium uptake between unbound and bound states were observed for peptides represented by grey bars, while significant differences were observed for those represented by light blue bars for MenB (**A**) and light-yellow bars (**B**) for Gc.

**Fig. S8.**
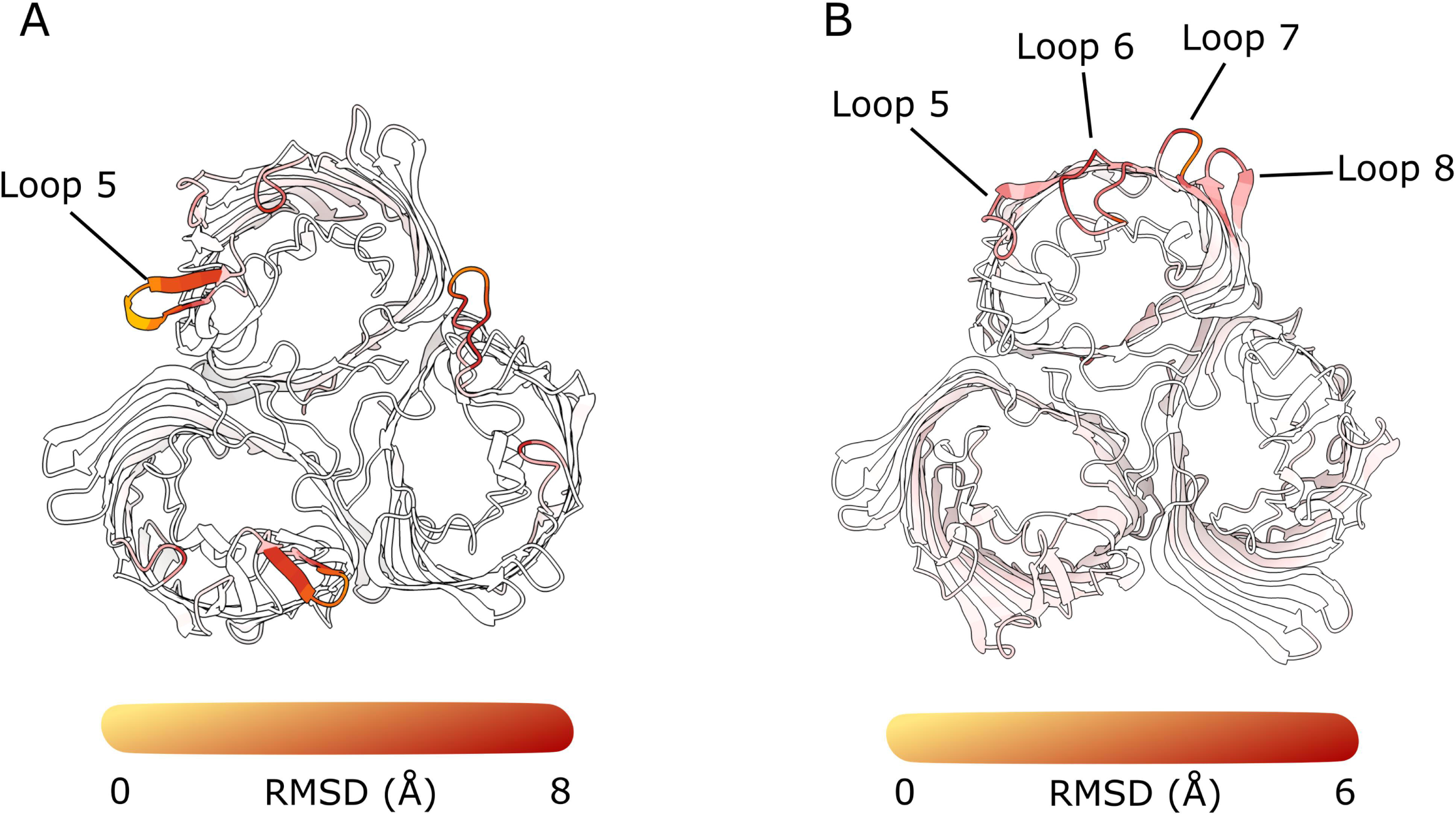
RMSD between PorB AF2 modeled orientations. (**A**) Gc-PorB. The RMSD has been calculated considering all the five models generated by AF2. The representative model has been colored-coded by RMSD showing high deviation in the region relative to loop 5. The RMSD range underlines conformational changes that could impair the monomer-monomer interface. (**B**) Men-PorB. The Root Mean Square Deviation (RMSD) has been calculated considering all the five models generated by AF2. The representative model has been colored-coded by RMSD showing high deviation in the region relative to loops 5, 6, 7, and 8. The RMSD range underlines conformational changes that do not involve the monomer-monomer interface.

## SUPPLEMENTARY TABLES

**Table S1.** Functional activity of 17 identified b-mAbs on Gc and MenB strains.

| Antigen recognized | mAb ID | Gc FA1090 |  | MenB strains – Bactericidal activity IC<sub>50</sub> (µg/ml) |  |  |  |  |  |  |  |  |  |
| --- | --- | --- | --- | --- | --- | --- | --- | --- | --- | --- | --- | --- | --- |
|  |  | Bactericidal activity (IC<sub>50</sub> µg/ml) | vOPA (%) | NZ98/254 | CU385 | H44/76 | MC58 | 5/99 | NGH38 | M07576 | ARG3191 | M01-240355 | M01-240364 |
| LOS | 01B05 | 2.03 | 50 | 0.98 | 0.24 | 0.24 | 0.12 | 0.24 | NB | 0.06 | NB | 7.81 | 0.98 |
| LOS | 03N18 | 4.00 | 40 | 0.49 | 0.24 | 0.24 | 0.06 | 0.49 | NB | 0.02 | 31.25 | 15.63 | 0.24 |
| LOS | 04E05 | 47.50 | 30 | 3.9 | 0.98 | 1.95 | 0.24 | 0.24 | NB | 0.03 | NB | 62.5 | 1.95 |
| LOS | 02G11 | 65.00 | 40 | 15.6 | 15.63 | 15.63 | 7.8 | 15.63 | 15.63 | 0.06 | NB | NB | 31.25 |
| PorB | 01H12 | 65.00 | 60 | 14 | NB | 125 | 62.5 | NB | NB | 112.5 | 0.12 | NB | NB |
| PorB | 02C05 | 65.00 | 60 | NB | NB | NB | NB | NB | NB | 0.12 | 0.06 | NB | NB |
| PorB | 02K03 | 40.30 | 37 | 7.8 | 62.5 | 125 | NB | NB | NB | 0.01 | 0.02 | NB | NB |
| PorB | 01J23 | 1.95 | 80 | 31.25 | 62.5 | 7.81 | 1.95 | NB | NB | 0.06 | 0.03 | 125 | NB |
| PorB | 01K12 | 0.002 | 80 | 3.9 | 15.63 | 0.49 | 0.24 | NB | NB | 0.02 | 0.01 | 1.95 | 125 |
| PorB | 03M18 | 1.95 | 50 | 62.5 | 62.5 | 15.63 | 3.91 | NB | NB | 0.06 | 0.03 | NB | NB |
| PorB | 04L22 | 0.004 | 0 | 0.98 | 3.91 | 0.49 | 0.49 | NB | 62.5 | 0.03 | 0.01 | 3.91 | NB |
| PorB | 01C05 | 0.006 | 80 | 7.8 | 7.81 | 0.49 | 0.49 | NB | NB | 0.02 | 0.01 | 0.98 | NB |
| PorB | 02G22 | 0.004 | 80 | 31.25 | 31.25 | 0.98 | 0.49 | NB | NB | 0.12 | 0.01 | NB | NB |
| Unknown | 01L10 | 65.00 | 40 | 31.25 | 62.5 | 125 | NB | NB | NB | 15.62 | 31.25 | NB | NB |
| Unknown | 01I20 | 1.00 | 70 | 31.25 | 15.63 | 0.49 | 0.98 | 125 | NB | 0.98 | 0.01 | 125 | NB |
| Unknown | 02E13 | 15.60 | 50 | NB | 31.25 | NB | NB | NB | NB | NB | 1.9 | NB | NB |
| Unknown | 05G16 | 125.00 | 30 | 62.5 | 31.25 | 15.63 | 31.25 | NB | NB | 31.25 | 0.03 | NB | NB |

**NB=** Not bactericidal

**Table S2.**
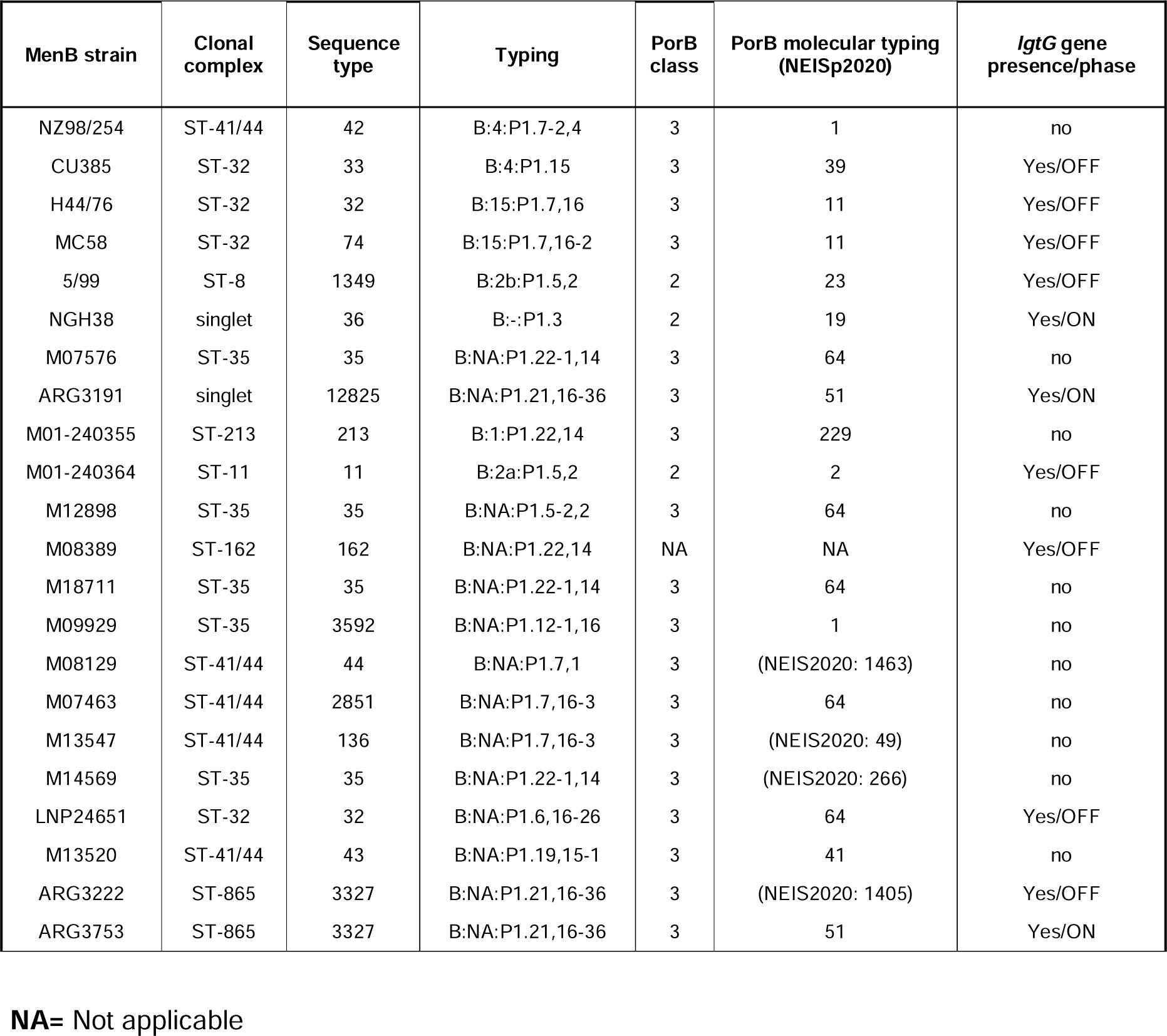
Summary of MenB strains used in the classical bactericidal assay and microarray.

| MenB strain | Clonal complex | Sequence type | Typing | PorB class | PorB molecular typing (NEISp2020) | *igtG* gene presence/phase |
| --- | --- | --- | --- | --- | --- | --- |
| NZ98/254 | ST-41/44 | 42 | B:4:P1.7-2,4 | 3 | 1 | no |
| CU385 | ST-32 | 33 | B:4:P1.15 | 3 | 39 | Yes/OFF |
| H44/76 | ST-32 | 32 | B:15:P1.7,16 | 3 | 11 | Yes/OFF |
| MC58 | ST-32 | 74 | B:15:P1.7,16-2 | 3 | 11 | Yes/OFF |
| 5/99 | ST-8 | 1349 | B:2b:P1.5,2 | 2 | 23 | Yes/OFF |
| NGH38 | singlet | 36 | B:-:P1.3 | 2 | 19 | Yes/ON |
| M07576 | ST-35 | 35 | B:NA:P1.22-1,14 | 3 | 64 | no |
| ARG3191 | singlet | 12825 | B:NA:P1.21,16-36 | 3 | 51 | Yes/ON |
| M01-240355 | ST-213 | 213 | B:1:P1.22,14 | 3 | 229 | no |
| M01-240364 | ST-11 | 11 | B:2a:P1.5,2 | 2 | 2 | Yes/OFF |
| M12898 | ST-35 | 35 | B:NA:P1.5-2,2 | 3 | 64 | no |
| M08389 | ST-162 | 162 | B:NA:P1.22,14 | NA | NA | Yes/OFF |
| M18711 | ST-35 | 35 | B:NA:P1.22-1,14 | 3 | 64 | no |
| M09929 | ST-35 | 3592 | B:NA:P1.12-1,16 | 3 | 1 | no |
| M08129 | ST-41/44 | 44 | B:NA:P1.7,1 | 3 | (NEIS2020: 1463) | no |
| M07463 | ST-41/44 | 2851 | B:NA:P1.7,16-3 | 3 | 64 | no |
| M13547 | ST-41/44 | 136 | B:NA:P1.7,16-3 | 3 | (NEIS2020: 49) | no |
| M14569 | ST-35 | 35 | B:NA:P1.22-1,14 | 3 | (NEIS2020: 266) | no |
| LNP24651 | ST-32 | 32 | B:NA:P1.6,16-26 | 3 | 64 | Yes/OFF |
| M13520 | ST-41/44 | 43 | B:NA:P1.19,15-1 | 3 | 41 | no |
| ARG3222 | ST-865 | 3327 | B:NA:P1.21,16-36 | 3 | (NEIS2020: 1405) | Yes/OFF |
| ARG3753 | ST-865 | 3327 | B:NA:P1.21,16-36 | 3 | 51 | Yes/ON |

**NA=** Not applicable

**Table 3.** Genetic characteristics of identified b-mAbs.

| Antigen | mAb ID | Clone ID | VH-JH usage | VH Mutation Freq (%) | HCDR3 Length (aa) | VL-JL usage | VL Mutation Freq (%) | LCDR3 Length (aa) |
| --- | --- | --- | --- | --- | --- | --- | --- | --- |
| LOS | 01B05 | 1 | IGHV2-5;<br>IGHJ4-1 | 12.29 | 15 | IGKV1-9;<br>IGKJ4-1 | 9.19 | 9 |
| LOS | 03N18 | 1 | IGHV2-5;<br>IGHJ4-1 | 24.22 | 15 | IGKV1-9;<br>IGKJ4-1 | 6.99 | 9 |
| LOS | 04E05 | 1 | IGHV2-5;<br>IGHJ4-1 | 22.05 | 15 | IGKV1-5;<br>IGKJ4-1 | 30.26 | 9 |
| LOS | 02G11 | 2 | IGHV1-2;<br>IGHJ3-1 | 25.84 | 13 | IGLV10-54;<br>IGLJ3-1 | 19.16 | 11 |
| PorB | 01H12 | 3 | IGHV4-34;<br>IGHJ4-1 | 2.37 | 24 | IGKV1-39;<br>IGKJ4-1 | 28.04 | 9 |
| PorB | 02C05 | 4 | IGHV4-34;<br>IGHJ4-1 | 8.14 | 28 | IGKV3-20;<br>IGKJ1-1 | 5.42 | 9 |
| PorB | 02K03 | 5 | IGHV4-34;<br>IGHJ4-1 | 4.41 | 22 | IGLV2-8;<br>IGLJ2-1 | 3.57 | 10 |
| PorB | 01J23 | 6 | IGHV4-34;<br>IGHJ6-1 | 8.14 | 22 | IGKV3-20;<br>IGKJ1-1 | 3.97 | 9 |
| PorB | 01K12 | 7 | IGHV4-34;<br>IGHJ6-1 | 8.47 | 23 | IGKV3-20;<br>IGKJ2-1 | 6.14 | 9 |
| PorB | 03M18 | 6 | IGHV4-34;<br>IGHJ6-1 | 8.47 | 22 | IGKV3-20;<br>IGKJ1-1 | 3.61 | 9 |
| PorB | 04L22 | 8 | IGHV4-34;<br>IGHJ3-1 | 7.80 | 26 | IGKV3-20;<br>IGKJ1-1 | 3.97 | 9 |
| PorB | 01C05 | 8 | IGHV4-34;<br>IGHJ3-1 | 8.81 | 26 | IGKV3-20;<br>IGKJ1-1 | 5.05 | 9 |
| PorB | 02G22 | 9 | IGHV4-34;<br>IGHJ3-1 | 9.15 | 23 | IGKV3-20;<br>IGKJ2-1 | 2.89 | 9 |
| Unknown | 01L10 | 10 | IGHV4-39;<br>IGHJ3-1 | 30.56 | 28 | IGKV3-20;<br>IGKJ3-1 | 6.50 | 10 |
| Unknown | 01I20 | 11 | IGHV4-34;<br>IGHJ3-1 | 8.47 | 25 | IGKV3-20;<br>IGKJ1-1 | 2.53 | 9 |
| Unknown | 02E13 | 12 | IGHV4-34;<br>IGHJ6-1 | 8.47 | 28 | IGKV3-20;<br>IGKJ4-1 | 3.61 | 9 |
| Unknown | 05G16 | 13 | IGHV4-34;<br>IGHJ4-1 | 7.46 | 23 | IGKV3-20;<br>IGKJ1-1 | 3.61 | 9 |

**Table S4.** Docking energy of 01K12/Gc FA1090 PorB complex. The docking energy reported for the lowest pose obtained for each Gc FA1090 PorB orientation.

| mAb ID | Orient-1 | Orient-2 | Orient-3 | Orient-4 |
| --- | --- | --- | --- | --- |
| 01K12 | -33.06396 | -47.7516 | -49.3324 | -39.50308 |

**Table S5.** Docking energy of 01K12/MenB NZ98/254 PorB complex. The docking energy reported for the lowest pose obtained for each MenB NZ98/254 PorB orientation.

| mAb ID | Orient-1 | Orient-2 | Orient-3 |
| --- | --- | --- | --- |
| 01K12 | -37.66491 | -42.57284 | -14.90061 |

## RESOURCE AVAILABILITY

### Lead Contacts

Further information and requests for resources and reagents should be directed to and will be fulfilled by the Lead Contacts, Oretta Finco, Emanuele Andreano and Rino Rappuoli.

### Materials Availability

Reasonable amounts of antibodies will be made available by the Lead Contacts upon request under a Material Transfer Agreement (MTA) for non-commercial usage.

## EXPERIMENTAL MODELS AND SUBJECT DETAILS

### Human samples ethics statement

Azienda Ospedaliera Universitaria Senese, Siena (IT) provided samples from 4CMenB vaccinees donors, who gave their written consent. The study was approved by the local Ethics Committee (AOU Senese, Parere nr. 13946_2018) and conducted according to good clinical practice in accordance with the declaration of Helsinki (European Council 2001, US Code of Federal Regulations, ICH 1997). Human complement source used in the study was obtained according to Good Clinical Practice in accordance with the declaration of Helsinki. Patients have given their written consent for the use of samples of study MENB REC 2ND GEN-074 (V72_92). The study was approved by the Western Institutional Review Board (WIRB).

### Animal model ethics statement

Animal husbandry and experimental procedures were ethically reviewed and carried out in accordance with European Directive 2010/63/EU, Italian Decree 26/2014 and GSK Vaccines’ Policy on the Care, Welfare and Treatment of Animals, and were approved by the Italian Ministry of Health (authorization 386/2021-PR). Upon arrival, animals were randomly distributed in different experimental groups in individually ventilated cages (IVC, Sealsafe Plus GM500 by Tecniplast). The acclimation lasted for a period of 5 days. At the end of the acclimation period, each animal was identified by an individual tattoo. All animals had ad libitum access to GMP-grade food (Mucedola 4RF25 TOP CERTIFICATE) and bottled, filtered, tap water. Certified, irradiated cellulose bags containing Mucedola SCOBIS UNO bedding, and carboard tunnels (ANTRUM) or plexiglass mouse house were provided within the cages. A few food pellets in the cage were also used as enrichment for forging and additional gnawing. Cage and bedding changes were performed once every week in according to protocol requirements. Air supplied in IVC will be 100% fresh air filtered by EPA filter by the IVC system, with 60-75 air changes per h. The animal room conditions were as follows: temperature 21°C (+/-3°C), relative humidity 50% (range 30-70%) and 12h/12h light/dark cycle. Pressure, temperature and relative humidity were recorded continuously by room probes, while the IVC system recorded individual motors’ performance. The light cycle setting was ensured by a qualified, alarmed system.

## MATERIALS AND METHODS

### 4CMenB OMVs labelling with Alexa488

4CMenB OMVs were fluorescently labelled targeting lysine residues with Alexa Fluor 488 (Alexa Fluor 488 carboxylic acid, succinimidyl ester, Invitrogen, A20000). OMVs were concentrated to 20 mg/ml in PBS 1× pH 7.2, following addition of AF488 dye in a 1:20 w/w ratio. After 18 h at room temperature (RT) in the dark with stirring, samples were purified through ultrafiltration with Amicon Ultra-4 100K using PBS 1× pH 7.2 for 30 washes. Labelled OMVs were characterized by SE-HPLC (TSK gel 6000PW column). Fluorescence intensities were measured at 490/525 nm Ex/Em, where an increased signal intensity was observed confirming the successful labelling, while absence of unconjugated dye was also verified. OMVs content was estimated through the Lowry method following manufacturing instructions (Pierce Modified Lowry Protein Assay Kit, Thermo).

### Single cell sorting of 4CMenB OMVs^+^ Memory B cells from vaccinees

Peripheral blood mononuclear cells isolation (PBMCs) and Single-cell Sorting were performed as previously described (Andreano et al., 2021; Huang et al., 2013). Briefly, PBMCs were isolated from heparin-treated whole blood by density gradient centrifugation (Ficoll-Paque PREMIUM, Sigma-Aldrich) and stained at RT for 20 min with Live/Dead Fixable Aqua Dead Cell Stain Kit (Invitrogen; ThermoFisher; cat#L34957) diluted 1:500. After incubation, cells were washed in Dulbecco’s modification of PBS (PBSB) and incubated with 20% of normal rabbit serum (Life Technologies) for 20Umin at 4U°C. PBMCs were then washed with PBSB and stained with 4CMenB OMVs labelled with Alexa488. Following 30Umin of incubation at 4U°C, cells were incubated with CD19 BV421 (BD, cat#562440), IgM PerCP-Cy5.5 (BD; cat#561285), CD27 PE (BD; cat#340425), IgDAlexa Fluor 700 (BD; cat#561302), CD3 PE-Cy7 (BioLegend; cat#300420), CD14 PE-Cy7 (BioLegend; cat#301814) and CD56 PE-Cy7 (BioLegend; cat#318318) at 4U°C. After incubation, stained PBMCs were single-cell-sorted using a BD FACSAria Fusion (BD Biosciences) in 384-well plates previously coated with 3T3-msCD40L feeder cells. Then, 4CMenB OMVs^+^ Memory B cells sorted were incubated for 10-14 days with IL-2 and IL-21 as previously described (Andreano et al., 2021; Huang et al., 2013).

### Whole-bacterial cell enzyme-linked immunosorbent assay (ELISA)

Bacteria were grown until mid-log phase as described and centrifuged at 4,500 x g for 5 min. Bacteria were resuspended in the same volume with PBS and seeded onto 384-well plates in a final volume of 20 µL. Incubation at 37°C, 5% CO_2_ for 30 min followed. Bacteria were fixed with 0.5% formaldehyde at RT for 30 min and then wells were washed twice with a washer dispenser (BioTek EL406, Agilent Technologies, US) with PBS, Tween20 0.05%, 150 µL/well/wash. Wells were washed and saturation step followed using PBS, BSA 1% in 50 µL to avoid unspecific binding and plates were incubated at 37°C for 1 h. After incubation, wells were washed, and primary antibodies contained into the TAP-supernatants were added in a 1:5 ratio in PBS, BSA 1%, Tween20 0.05% in 25 µL/well final volume and incubated for 1 h at 37°C without CO_2_. Wells were washed and 25 µL/well of alkaline phosphatase-conjugated goat anti-human IgG (Sigma-Aldrich, US) and IgA (Southern Biotech) were used as secondary antibodies. After 1h, a final wash followed and then pNPP (p-nitrophenyl phosphate) (Sigma-Aldrich) was used as substrate to detect the binding of the mAbs. The final reaction was measured by using the Varioskan Lux Reader (Thermo Fisher Scientific, US) at a wavelength of 405 nm. Samples were considered as positive if OD at 405 nm (OD_405_) was three times the blank.

### Single cell RT-PCR and Ig gene amplification

From the original 384-well sorting plate, 5 µL of cell lysate was used to perform RT-PCR and two round of PCRs as previously described (Andreano et al., 2021). Total RNA from single cells was reverse transcribed in 25 µL of reaction volume composed by 1 µL of random hexamer primers (50 ng/mL), 1 µL of dNTP-Mix (10 mM), 2 µL 0.1 M DTT, 40 U/mL RNase OUT, MgCl2 (25 mM), 5× FS buffer and Superscript IV reverse transcriptase (Invitrogen). Final volume was reached by adding nuclease-free water (DEPC). Reverse transcription (RT) reaction was performed at 42°C/10 min, 25°C/10 min, 50°C/ 60 min and 94°C/5 min. Heavy (VH) and light (VL) chain amplicons were obtained via two rounds of PCRs. All PCR reactions were performed in nuclease-free water in a total volume of 25 µL well. Briefly, 4 µL of cDNA were used for the first round of PCR (PCR I). PCR I master mix contained 10 mM of VH and 10 mM VL primer-mix, 10mM dNTP mix, 0.125 µL of Kapa Long Range Polymerase (Sigma), 1.5 µL MgCl2 and 5 µL of 5× Kapa Long Range Buffer. PCR I reaction was performed at 95°C/3 min, 5 cycles at 95°C/30 sec, 57°C/30 sec, 72°C/ 30 sec and 30 cycles at 95°C/30 sec, 60°C/30 sec, 72°C/30 sec and a final extension of 72°C/2 min. All nested PCR reactions (PCR II) were performed using 3,5 µL of unpurified PCR I product using the same cycling conditions. PCR II products were then purified by Millipore MultiScreen PCR 96 plate according to the manufacturer’s instructions. Samples were eluted in 30 µL nuclease-free water into 96-well plates and quantified by Qubit Fluorometric Quantitation assay (Invitrogen).

### Cloning of variable region genes and recombinant antibody expression in transcriptionally active PCR fragments

Vector digestions were carried out with the respective restriction enzymes AgeI, SalI and Xho as previously described by Tiller and colleagues. Briefly, 75 ng of IgH, Igλ and IgΚ purified PCR II products were ligated by using the Gibson Assembly NEB into 25 ng of respective human IgG1, IgΚ and Igλ expression vectors. The reaction was performed in 5 µL total volume. The ligation product was 10-fold diluted in nuclease-free water and used as the template for transcriptionally active PCR (TAP) reaction which allowed the direct use of linear DNA fragments for *in vitro* expression. The entire process consists of one PCR amplification step using primers that include functional promoter (human CMV) and terminator sequences (SV40) of the expression vectors onto the PCR II products. TAP reaction was performed in a total volume of 25 µL using 0.12 µL of Q5 polymerase (NEB), 5 µL of GC Enhancer (NEB), 5 µL of 5X buffer, 10 mM dNTPs, 0.125 µL of forward/reverse primers and 3 µL of ligation product. TAP reaction was performed by using the following cycling conditions: 98°C/2 min, 35 cycles 98°C/10 sec, 61°C/20 sec, 72°C/1 min and 72°C/5 min as final extension step. TAP products were purified under the same PCR II conditions, quantified by Qubit Fluorometric Quantitation assay (Invitrogen) and used for transient transfection in Expi293F cell line (Thermo Fisher Scientific) according to the manufacturer’s instructions.

### Gc and MenB strains

FA1090 and F62 Gc strains were purchased from American Type Culture Collection (ATCC). The authors are grateful to Dr. Darryl Hill of the University of Bristol (United Kingdom) for providing BG strains and to Dr. Sanjay Ram (University of Massachusetts, USA) for providing Gc MS11 LOS *lgt* mutants (Ram et al., 2018). The MS11 LOS *lgt* mutant strains were created in the background of Gc MS11 4/3/1, a variant of MS11 VD300 with an isopropyl-D-thiogalactopyranoside (IPTG)– inducible pilE that controls pilus expression. In these mutant strains the expression of the four phase-variable lgt genes (*lgtG, lgtA, lgtC* and *lgtD*) was genetically fixed either ON or OFF (or deleted). For kindly providing meningococcal serogroup B strains, the authors are grateful to: Xin Wang, Henju MarjukiXin Wang and Jarad Schiffer (CDC, Centers for Disease Control and Prevention, Atlanta, GA, USA) for M07576, M12898, M08389, M18711, M09929, M08129, M07463, M13547, M14569 and M13520; M-K. Taha (Institut Pasteur, Paris) for LNP24651 strain, Richard Moxon (University of Oxford, Oxford, United Kingdom) for MC58; Diana R. Martin (Institute of Environmental Science and Research, Porirua, New Zealand) for NZ98/254; Ray Borrow (Health Security Agency, Manchester, United Kingdom) for M01-240355 and M01-240364; Dominique A. Caugant (NIPH, Norwegian Institute of Public Health, Oslo, Norway) for H44/76, NGH38, CU385 and 5/99; Adriana Efron (Instituto Nacional de Enfermedades Infecciosas-ANLIS “Dr. Carlos G. Malbrán”, Buenos Aires, Argentina) on behalf of the Argentinian National Laboratories Network (NLR) for ARG3191, ARG322 and ARG3753. For kindly providing Gc low passage clinical isolates the authors are grateful to Darryl Hill (University of Bristol).

### Bacterial genome sequencing

For clinical isolates reported in Table S2, Neisserial genomic DNA was extracted from cell suspensions of overnight culture plates using the GenElute Bacterial Genomic DNA Kit (Sigma). Genomic libraries were generated using Illumina Nextera DNA Flex Library Prep according to the manufacturer’s instructions and sequenced on Illumina MiSeq 2×250bp paired end platform. The read sequences from all the samples above were assembled using Spades, version 3.13, with default parameters. The resultant assemblies, in addition to public genomes downloaded from the PubMLST website were uploaded to the internal Neisseria PubMLST database that was used for the typing of our loci of interest. LOS beta chain presence or absence was inferred from the genetic information about *lgtG* locus. BLAST of *lgtG* sequence from MC58 and 1’000 flanking nucleotides were used to identify *lgtG* locus, if present. Then the *lgtG* sequence extracted was translated in order to understand whether the sequence could lead to a functional protein product or not. In case of absence of the *lgtG* locus or short protein translation, the LOS beta chain was predicted to be absent, in case of presence of a full length *lgtG* enzyme, compatible with a functional protein product, the LOS beta chain was predicted to be present.

### Bacterial growth

Fresh cultures of different Gc strains were prepared from frozen stocks by streaking onto gonococcal agar (GCA) consisting of agar base supplemented with 1% v/v IsoVitaleX (BD Biosciences, Franklin Lakes, NJ, USA). On the following day, bacteria were grown in gonococcal (GC) liquid medium at 37°C starting from OD_600_ 0.1 until mid-log phase cultures, i.e., OD_600_ 0.5

### mAb screening by Resazurin-based Antibody Bactericidal Assay (R-ABA)

The initial bactericidal screening on Gc was preformed through R-ABA as previously described^19^. Briefly, baby rabbit complement (BRC) (Cedarlane) was used as a source of complement in R-ABA assays. BRC was diluted to 10% v/v for FA1090 and F62 strains, while 20% v/v was used for BG27 strain. BRC was heat-inactivated (hiBRC) at 56°C for 30 min before use, as a complement inactivated control. Bacteria were grown to mid-log phase and resuspended in PBSB as detailed above. Reactions were performed by incubating bacteria with TAP supernatants in PBSB, 2% FBS and 0.1% glucose and in the presence of 10% v/v of BRC round bottom 96-well plates. hiBRC control adjusted to a final volume of 50 mL in PBSB, 2% FBS and 0.1% glucose was used. Reactions were incubated for 2 h at 37°C, 5% CO_2_. Then, 10 µL of 0.025% resazurin solution in sterile distilled water were added to each well. The reactions were then further incubated for 2 h at 37°C, 5% CO_2_. At the end of the incubation, fluorescence signals were measured by a Varioskan LUX multimode microplate reader (Thermo Fisher, Waltham, MA, USA) using 560 nm for excitation and 590 nm for emission. After the measurement, the assay plate was kept at 37°C ON to allow complete conversion of resazurin to resorufin in the wells containing live bacteria. On the following day, the plate was observed by eye and pictures were taken.

### Flask expression and purification of mAbs

Expi293F cells were transiently transfected with plasmids carrying the antibody heavy chain and the light chains with a 1:2 ratio, respectively. The transfections last for six days at 37°C with 8% CO_2_ in shaking conditions at 125 rpm according to the manufacturer’s protocol (Thermo Fisher Scientific, US). ExpiFectamine 293 transfection enhancers 1 and 2 were added 16 to 18 h post-transfection to boost cell viability and protein expression. Cell cultures were harvested six days after transfection. Supernatants collected were then pooled and clarified by centrifugation (4,500 x g, 15 min, 4°C) followed by filtration through a 0.22 mm filter. Protein A chromatography (HiTrap Protein A HP, Cytiva) was used for antibody capture from cell culture supernatant and purification. The supernatants, diluted 1:1 with buffer A (50 mM NaH2PO4, 300 mM NaCl, pH 8.0) were loaded onto the equilibrated protein A column with buffer A. After column washing with 10 bed volumes of buffer A, bound antibodies were eluted with buffer B (0,1 M citric acid, 300 mM NaCl, pH 3,6). The eluted antibodies were neutralized with 10% of the final volume per fraction of 1 M Tris-HCl buffer, pH 9. The fractions of interest identified by following the chromatographic profile (absorbance at 214 nm), were pooled and then buffer exchanged with O/N dialysis at 4°C against PBS buffer (≥100× the volume of the antibody solution). Monomeric state of purified mAbs was checked by analytical gel filtration (SE-UPLC) with the Superdex 200 Increase 5/150 GL (Cytiva) column by using PBS as running buffer (flow rate 0,4 ml/min) and by following absorbance at 215 nm. MAbs with a purity degree less than 85% were subjected to an additional chromatographic step using a preparative Superdex 200 column (GE) in PBS buffer in order to remove aggregates or degradations. Then additional analytical SEC were performed to assess the final level of purity of purified mAbs. After a sterile filtration with 0,22 µm filter, the concentration of final batch was determined by absorption at 280 nm by using a NanoDrop 8000 spectrophotometer (Thermo Scientific). Final mAb samples were also analyzed by SDS-Page gel in order to compare migration of reduced/denatured and not reduced/not denatured samples using the NuPage gel system (Invitrogen) with MES as running buffer. Gels were stained with ProBlue safe stain (Giotto Biotech). The mAbs were then stored at -80°C.

### LOS extraction from 4CMenB OMVs

LOS was extracted using a general method based on the Westphal hot phenol extraction process to purify whole LOS from 4CMenB OMVs samples (Westphal, 1965). Briefly, OMVs were stirred at 65°C until the temperature equilibrated. An equal volume of 90% (w/v) phenol which had been preheated to 65°C was added and thoroughly mixed for 30 min. The resulting mixture was rapidly cooled by stirring for 30 min in an ice-water bath. The phenol mixture was then centrifuged at 4°C at 4,000 x g for 10 min. A sharp interface occurred between the aqueous, phenol, and interface layers. The aqueous and phenol layers were removed by aspiration. The aqueous layer containing the lipopolysaccharide was retained while the phenol layer was discarded. Cold ethanol precipitation was performed 3-4 times on the aqueous phase and the final pellet was suspended in distilled water. The sample was then mixed with a solution of Proteinase K–Agarose from *Tritirachium album* (Sigma) and incubated for 16 h at RT with stirring. The sample mixture was then centrifuged at 15,000 x g for 10 min and supernatant containing LOS was collected and separated from the agarose pellet. The LOS sample was finally purified through ultrafiltration with Amicon Ultra-4 10K.

### LOS quantification

The content of LOS extracted or present in the OMV samples was determined by the quantification of the reactive carbonyl groups of the saccharide moiety, generated after acid hydrolysis to remove the Lipid A and derivatized with semicarbazide (SCA), by high-performance liquid chromatography–size exclusion chromatography (SE-HPLC) analysis. The SCA reaction coupled with SE-HPLC analysis has been already reported in literature (Micoli et al., 2014). In the first step, samples (extracted LOS or OMV) were treated with acetic acid (1% final concentration) and hydrolyzed for 2 h at 100°C to remove the Lipid A. After the hydrolysis, each sample was dried in a SpeedVac system to remove the acetic acid, and then dissolved with MilliQ water. Samples were centrifuged for 10 min at 15,000 x g to separate pellet (Lipid A) and supernatant (OS). To obtain UV detectable samples, the supernatants were derivatized with semicarbazide (SCA). A stock SCA solution was prepared dissolving 100 mg of SCA hydrochloride and 90.5 mg of sodium acetate in 10 mL of MilliQ water. Equal volumes of sample and SCA solution were transferred into clean vials (i.e., 100µL sample + 100µL SCA solution) and incubated in a pre-heated water bath at 50°C for 50 min. Samples were chilled at 2-8°C for 15 min and then filtered in HPLC vials. The KDO content of the GMMA samples is quantified based on a calibration curve prepared with standard KDO ammonium salt solution. The LOS content is expressed in nmol/mL of KDO, which matches the nmol/mL of OS.

### Immunoblot analysis of mAbs on extracted LOS

Extracted LOS was loaded on SDS-PAGE gel at 0.008 nmol/well. OMV samples were normalized by LOS quantity obtained from semicarbazide derivatization/SE-HPLC method. Samples were run on a 16% Tris-glycine SDS-PAGE gel using a Tris-glycine 1× buffer. The ladder consisted of the following proteins: Bradykinin (1,060 Da), Insulin Chain B (3,496 Da), Aprotinin (6,500 Da), α-Lactalbumin (14,200 Da), Myoglobin (17,000 Da) and Triosephosphate Isomerase (26,600 Da). Bands were transferred to nitrocellulose membranes (The iBlot Kit Thermofisher) and membranes were blocked with PBS 1× + BSA 3% + Tween20 0.05% for 1 h at RT. mAb concentration was normalize to 1μg/mL in PBS 1× + Tween20 0.05% and incubated for 1 h at RT. Signals were visualized with anti-human IgG alkaline phosphatase (Sigma, diluted 1:2,000 in PBS 1× + Tween20 0.05%) incubated for 30 min at RT, followed by AP Conjugate Substrate kit (Biorad) for 5 min at RT.

### Immunoblot on 4CMenB OMVs

This mixture was run on an SDS-PAGE gel (NuPAGE 4-12% Bis-Tris Gel) in MES Buffer 1X (NuPAGE MES SDS Running Buffer). Samples were transferred on a nitrocellulose membrane (iBlot 2 NC, Thermo Fisher) using an Invitrogen iBlot 2 Gel Transfer Device. After transfer, the membrane was blocked with 5% non-fat dry milk in 20 mM TBS, 0.05% tween-20. After blocking, the membrane was incubated with 2 ug/ml of antibody of interest in 5% milk in 20 mM TBS, 0.05% tween-20 ON at 4 °C. The membrane was then washed 3 times with 20 mM TBS, 0.05% Tween 20 (five min per wash) and then incubated with the secondary antibody (secondary antibody anti-human Fab 1:100,000) in 5% milk in 20 mM TBS, 0.05% tween-20 for 1 h at 4 °C. After 3 washes in 20 mM TBS, 0.05% Tween-20, the membrane was developed using Thermo Scientific™ Pierce™ ECL Western Blotting Substrate and imaged using the Invitrogen iBright Imaging system with the chemiluminescence detection method.

### OMV preparation

To produce OMVs, Gc and MenB strains were plated on GC +1% Isovitalex or GC agar plates, respectively. Plates were incubated ON at 37°C in 5% CO_2_. The following day, MenB colonies were inoculated in 10 mL of Mueller-Hinton Broth (OD of U0.05) and allowed to grow with shaking until OD of U1.0-1.5 at 37°C. Then 10 mL were put in 50 mL of prewarmed slightly modified MCDMI medium and incubated at 37°C in 5% CO_2_. OD_600_ was constantly monitored, and the growth was stopped when OD_600_ remained stable for 1 h and 30 min. Gc colonies were instead inoculated in 5 ml of GC +1% Isovitalex and the growth was followed for 28 h in 24 deep-well plate at 37°C 350 rpm. Bacteria cultures were collected and discarded by centrifugation for 60 min at 4,000-8,000 x g and the supernatants were subjected to high-speed centrifugation at 11,9000 x g for 2-3 h at 4°C (Beckman Coulter Optima Ultracentrifuge). The pellets containing the OMVs were washed with PBS, ultracentrifuged again, as above described, and finally resuspended in PBS. OMV total protein content was quantified through the Lowry assay (DC Protein Assay, BioRad) following manufacturer’s instructions.

### Protein array design, generation, validation and hybridization

Monoclonal Abs were tested over two separate protein microarrays previously generated (Viviani et al., 2023). Specifically, the recombinant protein microarray, encompassed 12 recombinant proteins and the three recombinant meningococcal antigens of the 4CMenB vaccine (NHBA-GNA1030; GNA2091-fHbp and NadA) spotted at 0.5Umg/mL in 40% glycerol, while the vesicles protein chip containing 26 recombinant *E. coli* GMMAs, two GMMAs empty and OMVs from NZ98/254. The latter array was expanded with 14 different meningococcal OMV (0.5 mg/mL in 20% glycerol) and 18 different gonococcal OMVs (0.25Umg/mL in 20% glycerol). Controls consisted of 8 serial two-fold dilutions of human IgG (from 0.5Umg/mL to 0.004Umg/mL in 40% glycerol), unrelated proteins (0.5Umg/mL in 40% glycerol) and PBSU+U40% glycerol spots. Each sample was spotted randomly in replicates per array onto ultra-thin nitrocellulose coated glass slides (FAST slides; Maine Manufacturing). Printing was performed with the ink-jet spotter Marathon Argus (Arrayjet) (200 pl each spot) in a cabinet with controlled temperature and humidity (18U°C and 50–55%, respectively). To ensure efficient and reproducible protein immobilization a preliminary array validation was carried out with anti-FLAG antibodies (Sigma-Aldrich, cat# F7425) 1:5000 and mouse anti-His_6_ tag polyclonal antibodies (Thermo Fisher, cat# 37-2900) 1:1,000, followed by detection with an AlexaFluor 647-conjugated anti-rabbit or anti-mouse IgG secondary antibody (Jackson Immunoresearch, cat# 111-605-046, cat# 115-605-174) -1:800. Preliminary experiments with mAbs showed that 0.5 µg/mL corresponded to the best signal to noise ratio. For mAbs hybridization experiments, nonspecific binding was minimized by preincubating the slides with a blocking solution (BlockIt, ArrayIt) for 1Uh. mAbs were then diluted to 0.5 µg/mL in BlockIt buffer and overlaid for 1Uh at RT prior to undergo two washes with Tween 0.1% in PBS (TPBS). AlexaFluor 647-conjugated anti-human IgG secondary antibody (Jackson Immunoresearch, cat# 115-605-174) diluted 1:800 was incubated 1 h, before proceeding with slide scanning. Fluorescence images were obtained using InnoScan 710 AL (Innopsys) and the images were generated with Mapix software at 10Uμm/pixel resolution. ImaGene 9.0 software (Biodiscovery Inc.) was used to calculate spot fluorescence intensities while the microarray data analysis step was carried out with an *in-house* developed R script. For each protein the Mean Fluorescence Intensity (MFI) of replicates was obtained after the subtraction of local background values surrounding each spot. MFI were greater than 6,000, corresponding to the MFI of control protein spots after detection with fluorescent-labelled antibodies, plus ten times the standard deviation, were considered positive. MFI scores were ranked in four categories: (1) high reactivity; MFIU≥U30,000; (2) medium reactivity; 15,000U≤UMFIU>U30,000; (3) low reactivity; 6,000U≤UMFIU>U15,000; (4) no reactivity; MFIU<U6,000.

### Killing-based Serum Bactericidal Assay (SBA) on FA1090 Gc strain

Functional characterization of b-mAbs was performed by classical serum bactericidal assay (SBA) against FA1090 strain. Human serum obtained from volunteer donors with no detectable intrinsic bactericidal activity was used as source of complement. Bacteria were grown at 37U°C in GC liquid medium supplemented with 1% Isovitalex until mid-exponential phase (OD_600_ 0.5). FA1090 growth was also supplemented with 1 µg/mL of CMP-NANA (Cytidine-5′-MonoPhospho-N-Acetyl NeurAminic acid sodium salt). Then, bacteria were diluted in DPBS, with 0.1% glucose and 1% BSA to a working dilution of 10×10^3^UCFU/mL. Subsequently, bacteria were two-fold diluted and incubated with 10% human complement and mAbs for 1h at 37°C. After the incubation, 100 µL of GC medium plus 0.5% of Bacto Agar, was added to the reaction mixture and incubated O/N at 37C° with 5% CO_2_. The day after, the plate well images were automatically acquired with a high throughput image analysis system and the Colony Forming Units (CFUs) were automatically counted for each well by an internal customized colony counting software. Bactericidal titer was defined as 50% decrease in CFU number compared to the reaction mixture without antibody.

### Killing-based Serum Bactericidal Assay (SBA) on MenB strains

Bactericidal activity of mAbs to MenB strains was carried out through classical SBA in the presence of 25% baby rabbit complement (Cedarlane). The reaction was performed in 96-well plate. From the glycerol stock, bacteria were seeded and grown O/N on chocolate agar plate at 37 °C in 5% CO_2_. The day after 10-15 colonies were inoculated in Müller-Hinton broth containing 0.25% glucose to reach an OD_600_ of 0.05 to 0.06. Bacteria were then incubated at 37 °C shaking until OD_600_ 0.25 (U10^9^ colony forming units CFU/ml). Bacteria were diluted 30,000-fold in DPBS, 1% (w/v) BSA, 0.1% glucose (w/v) and added to a reaction mix with a two-fold serial dilution of monoclonal antibody and baby rabbit complement. The plate was incubated for 1 h on shaking at 37 °C in 5% CO_2_. After 1 h incubation 100µL of melted TSB + 0.7% agar medium was added in each well allowing for the solidification. Then a second layer of 50 µL of melted agar medium was added in each well until the solidification. Plate was then incubated O/N at 37°C. The day after, the plate well images were automatically acquired and the bactericidal titer was calculated as for Gc strains, above indicated.

### Binding characterization by flow cytometry

All FACS experiments were carried out in the same conditions. After reaching OD_600_ 0.5, bacteria were centrifuged at 4,500 x g, 5 min and then resuspended and diluted in PBSB, 1% BSA to OD_600_ 0.2. 50 mL of bacterial suspension were seeded onto 96-well round bottom TC-treated microplates (Corning, US). Bacterial suspensions were centrifuge as described and resuspended in a mix with PBSB, 1% BSA and primary antibodies at 10 mg/mL in 50 mL. An incubation step of 1 h followed, at 37°C, 5% CO_2_. Bacterial suspensions were centrifuge and resuspended in a mix of PBSB, 1% BSA and a goat anti-Human IgG secondary antibody, labeled with Alexa Fluor 488 (Thermo Fisher Scientific, US) in 50 mL. Bacterial suspensions were centrifuge (4,500 x g, 5 min) and fixed with 0.5% formaldehyde at RT for 30 min. After fixation, bacterial suspensions were centrifuge (4,500 x g, 5 min) and re-suspended in PBS in a final volume of 50 mL. The samples were read using BD FACS Canto II flow cytometer. 10,000 counts were acquired for each sample. The analysis was carried out using FlowJo (software version 10).

### Immunofluorescence analyses on FA1090 Gc strain

Bacteria were grown and re-suspended in PBSB to reach OD_600_ 0.2. 50 mL of diluted bacterial suspensions were seeded onto a 96-well glass-bottom plates (Cell imaging plate, Eppendorf) and incubated for 30 min at 37°C, 5% CO_2_ to allow for bacteria adhesion. After incubation, bacteria were fixed for 30 min at RT with 0.5% formaldehyde. After two washing steps, 50 mL of PBSB, 1% BSA were added, and the plate was incubated for 1 h at 37°C. After saturation, a mixture containing PBSB, 1% BSA and primary antibodies (10 mg/mL) in 50 ml was added. Samples were incubated for 1 h at 37°C. A washing step with 100 mL of PBS followed the incubation. A goat anti-Human IgG secondary antibody labelled with Alexa Fluor 488 was added (Thermo Fisher Scientific). The plates were incubated for 30 min at 37°C. Following two washing steps as described, 4’,6-diamidino-2-phenylindole (DAPI) was prepared at 0.1 mg/mL in PBSB a final volume of 50 mL/well. Bacterial pellet was resuspended and incubated for 30 min at 4°. Z-stack images were acquired with a spinning disk super-resolution microscope (CSU-W1-SoRA Nikon) with a 60X oil objective (numerical aperture 1.49) and a Photometrics BSI sCMOS camera using the same settings. 3D Deconvolution (Blind method, 20 iterations) and denoise were applied to high-resolution images. 3D reconstructions were obtained using Fiji software (version 2.1.0). Total fluorescence intensity was quantified using Fiji from the sum intensity projection of the confocal *z*-stack images after segmentation using Otsu thresholding (Otsu, 1979).

### Immunogold analyses of FA1090 Gc strain

FA1090 strain was grown as already described until OD_600_ 0.5. After growth, 2.5 mL of bacterial suspension were fixed with 4% formaldehyde for 10 min, RT in a final volume of 5 mL. Then, samples were centrifuged at 3,000 x g for 7-10 min 25°C and resuspended in 5 mL of DPBS. Subsequently, 5 µL of fixed bacterial suspension were adsorbed to 300-mesh nickel grids and blocked in a mixture of PBS 1% BSA and mAbs (diluted 1:500 in PBS) for 1 h. Grids were washed several times with PBS and incubated with 12-nm gold-labeled anti-human secondary antibody (Jackson ImmunoResearch, diluted 1:40 in PBS) for 1 h. After several washes with distilled water, grids were air-dried. Images were acquired using a 120kV TEM FEI Tecnai G2 spirit microscope along with the Tvips TemCam.

### Visual opsonophagocytosis assay (vOPA)

THP-1 cells were seeded and differentiated into 96 well plates. After 5 days of differentiation, cells were infected with FA1090::sfGFP strain. Bacteria were grown until mid-logarithmic phase and pre-incubated with mAbs supernatants diluted 1:5 in RPMI media. After 30 min of pre-incubation, mAbs and bacteria mix were added onto dTHP-1. To synchronize the infection, the 96 well plates were centrifuged for 1 min at 200 x g. After 1h of infection, each well was fixed with 2% PFA for 15 min, blocked with 1% (w/v) BSA. Extracellular FA1090::sfGFP was stained with the primary antibody 2C7, at final concentration of 3 mg/ml for 1h at RT. Subsequently, a secondary antibody goat anti-Human IgG Alexa Fluor 568 (Thermo Fisher, A-21090) was added using a dilution factor of 1:2000 and incubated 30 min at RT. CellMask™ Deep Red stain (Invitrogen) was used to stain the membrane, providing a mean to delineate the cell boundary and DAPI to stain the nucleus. Images were automatically collected with microscope Opera Phenix High-Content Screening System (PerkinElmer) using an objective magnification of 40×, acquiring 16 fields of view and 13 z-stacks each per well. THP-1 cell detection was performed on DAPI (nuclei) and CellMask (cell membrane). While bacteria were segmented by colocalizing DAPI and GFP. Overlapping bacteria with CellMask were considered as internalized ones, and cells were considered as infected. The image analysis pipeline was performed using Harmony Software. The read-out (phagocytic activity) was determined on the total number of internalized bacteria / total number of infected cells.

### HDX-MS on MenB and Gc PorB

Epitope mapping of PorB antigen, embedded on deoxycholate extracted MenB and Gc OMVs with b-mAb 01K12, was performed by Hydrogen Deuterium eXchange associated to Mass Spectrometry (HDX-MS), comparing the amount of deuterium incorporated by PorB peptides in presence and absence of antibody. PorB amount in the dOMV was estimated to be 40 % of the total protein content as previously reported (Tani et al., 2014). The antigen alone (60pmol of OMVs-embedded PorB) or antibody/antigen mixture (OMVs-embedded PorB/mAb 1/2 molar ratio) were incubated for 30 minutes at 25°C. The labelling procedure was carried out in an ice bath and was initiated by adding deuterated PBS buffer (pD of 7.3), reaching a deuterium excess of more than 90% over five time points ranging from 15 seconds to 100 min (15 s, 1 min, 5 min, 30 min, 100 min) (Malito et al., 2013). Samples were quenched and delipidated by TCA precipitation and acetone washes as previously reported (Donnarumma et al., 2018). Experimental replicates for statistical analysis were prepared for both unbound and bound states for the 15sec D2O exposure. Samples were injected into a NanoAcquity UPLC with HDx technology (Waters Corporation, Milford, USA), digested on-line with a homemade pepsin column and the mass spectra of peptic fragments, desalted, and separated by reverse-phase ultraperformance liquid chromatography (RP-UPLC), were acquired in resolution mode (*m/z* 200–1200) on a SynaptG2Si mass spectrometer with a standard electrospray ionization source. Peptides were identified by MS^E^ analysis. Data were processed using Protein Lynx Global Server 3.0 and DynamX 3.0 software (Waters) was used to select peptides for the analysis. Only the peptic peptides present in at least four out of six repeated digestions of the unlabeled proteins, presenting at least 0.2 identified fragments per amino acid residues, were selected for the HDX-MS analysis. For statistical analysis three labeling reaction experiments of the antigen alone and in complex with the mAb were performed for the 15 seconds D2O exposure time point.

Analysis was performed by calculating a confidence interval (CI) based on the standard deviation (SD) in deuterium uptake at the time point performed in triplicates. For each state, the SDs were calculated using the root-mean-square as shown in following equation:

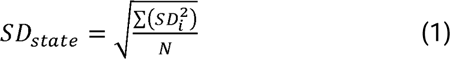

where *N* is the total amount of peptides measured in replicate. The pooled SD, for the two states, was calculated using the Equation (2):

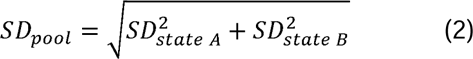

The pooled SD was utilized to identify the 98% CI through the equation (3):

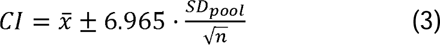

Where *x* is the assumed zero-centered average difference in deuterium uptake (*x*=0), 6.965 is the t-value corresponding to a two-tailed distribution with two degrees of freedom, and *n* s the analyzed sample size (set to 3 since the experiment was performed in triplicates).

### *In silico* docking of Gc PorB/01K12 and MenB PorB/01K12

The *in silico* docking experiments were conducted on the trimeric structural organization of MenB PorB (NZ98 strain) and Gc PorB (FA1090 strain) with version 2.4 of the HADDOCK software (van Zundert et al., 2016). The sequence alignment between the selected PorB strains and the publicly available PorB structures showed poor similarity. Therefore, molecular modeling was performed with a state-of-the-art approach namely AlphaFold 2 to provide a starting point for the docking analysis (Jumper et al., 2021). A total of five models for each strain, namely orient 1-5, were obtained and used for further investigation. Interestingly, the predicted Local Difference Test (pLDDT) ranged from approximately 70 in the loop regions, to 98 in the beta-barrel portion of the protein, highlighting (i) high model accuracy, and (ii) variability in the loop conformations. Loops 5-8 in MenB PorB and loops 4-7 in Gc PorB showed higher flexibility compared to the other modeled loops. Similarly, the variable region of the 01K12 antibody was modeled with DeepAb (Ruffolo et al., 2022), an artificial intelligence approach that provides a highly confident estimation of the CDR3. The epitope was identified in both strains with a total of eight regions: 39-46 (loop1), 81-95 (loop2), 113-143 (loop3), 164-176 (loop4), 197-210 (loop5), 234-249 (loop6), 270-284 (loop7) and 309-319 (loop8). The paratope region of the selected mAbs was identified with Paragraph (Chinery et al., 2023) and defined in HADDOCK as “active”, while the epitope regions on PorB as “passive”, meaning the paratope region needs to contact at least one of the PorB residues and there is no penalty if it does not contact all of them, allowing the mAb to freely explore the binding loops. All three docking iterations, it0, it1, and water, generated 5000, 400, and 400 poses, respectively, using the default values and scoring function. Clustering was performed based on backbone RMSD with a distance cut-off of 5 Å on the latest 200 generated poses. Finally, the lowest score was used to select the “best cluster” as the most antigen/antibody interaction representative.

### Mouse infection studies

Specific and Opportunistic Pathogen Free (6–8-week-old) female SOPF-BALB/c mice (The Charles River Laboratories, France) in the diestrus or anestrus stages of the estrous cycle received 21-day slow-release 17-β estradiol pellet (Innovative Research of America) implanted subcutaneously to induce susceptibility to gonococcal colonization. Antibiotics were also administered to suppress the overgrowth of commensal flora that occurs under estradiol treatment (Jerse, 1999). Vancomycin (4mg/mL; Sigma) and Streptomycin sulfate (24mg/ml; Sigma) were administered via intraperitoneal injection by the following dosing regimen: a single injection of 0.25 ml on day -2 and 0.15 ml given twice daily from day -1 to day +1. Trimethoprim (0.4 g/L; Sigma) was provided in the drinking water through day + 1 and both trimethoprim (0.4 g/L) and streptomycin sulfate (5g/L) were provided in the drinking water from day +2 through the remainder of the study period. Two days after pellet implantation, mice were inoculated intravaginally with 20 µl of FA1090 strain harvested from ON growth on GC agar plates supplemented with 1% v/v IsoVitaleX (BD Biosciences, Franklin Lakes, NJ, USA) and suspended in PBS (OD_600_=0.1) at a dose that establishes infection in 80-100% of mice (around 2×10^6^ CFU/mouse). Following inoculation, the vaginal mucosae of test mice were cultured each day by gently inserting a Dacron swab (PurFybr, Inc., Munster, Ind.) into the vagina. Gc FA1090 strain, used for the *in vivo* efficacy study, is a serum resistant PorB.1B, streptomycin resistant (SmR) strain originally isolated from a female with disseminated gonococcal infection (DGI) (Cohen and Cannon, 1999). The efficacy of mAb 01K12 was examined by daily intravaginal administration throughout the course of the experiment. To test mAb 01K12 specificity, additional groups of mice received intravaginally an unrelated a-RSV mAb (non-specific control) and saline (vehicle control). Saline and monoclonal antibodies treatment began 2 h prior to the challenge and were performed 1 h after vaginal sampling in the following days (Gulati et al., 2019b). Both a-RSV and 01K12 mAbs were injected at dosage of 0.5 µg since in previous studies a non-specific effect of anti-RSV mAb injected intravaginally in higher amounts (10 and 1 μg) was observed. As described above, gonococcal CFUs were enumerated by vaginal swabbing performed up to 7 days post challenge. Daily bacterial burdens were measured by first ringing vaginal swabs in 100 μL of saline and then plating serial 10-fold dilutions onto supplemented GC agar with VCNT (vancomycin, colistin, nystatin and trimethoprim sulfate; Becton Dickinson) and 100 µg/ml of streptomycin sulfate (Sigma). The limit of detection was 10 CFU per swab eluted in 100 μL saline. Analyses of the data are described below. Clearance of infection was defined by 3 or more consecutive days of negative cultures. Daily vaginal swabs were also cultured on Heart infusion agar (HIA; BD) agar plates to isolate facultatively anaerobic commensal bacteria. Incubation conditions for *N. gonorrhoeae* and for commensal bacteria were at 37°C in a humid atmosphere containing 5% CO_2_.

## REFERENCES

Abara, W.E., Bernstein, K.T., Lewis, F.M.T., Schillinger, J.A., Feemster, K., Pathela, P., Hariri, S., Islam, A., Eberhart, M., Cheng, I., et al. (2022). Effectiveness of a serogroup B outer membrane vesicle meningococcal vaccine against gonorrhoea: a retrospective observational study. The Lancet Infectious Diseases 22, 1021–1029.

Andreano, E., Nicastri, E., Paciello, I., Pileri, P., Manganaro, N., Piccini, G., Manenti, A., Pantano, E., Kabanova, A., Troisi, M., et al. (2021). Extremely potent human monoclonal antibodies from COVID-19 convalescent patients. Cell 184, 1821–1835.e1816.

Boslego, J.W., Tramont, E.C., Chung, R.C., McChesney, D.G., Ciak, J., Sadoff, J.C., Piziak, M.V., Brown, J.D., Brinton, C.C., Jr., Wood, S.W., et al. (1991). Efficacy trial of a parenteral gonococcal pilus vaccine in men. Vaccine 9, 154–162.

Bruxvoort, K.J., Lewnard, J.A., Chen, L.H., Tseng, H.F., Chang, J., Veltman, J., Marrazzo, J., and Qian, L. (2023). Prevention of Neisseria gonorrhoeae With Meningococcal B Vaccine: A Matched Cohort Study in Southern California. Clinical infectious diseases : an official publication of the Infectious Diseases Society of America 76, e1341–e1349.

Chakraborti, S., Lewis, L.A., Cox, A.D., St Michael, F., Li, J., Rice, P.A., and Ram, S. (2016). Phase-Variable Heptose I Glycan Extensions Modulate Efficacy of 2C7 Vaccine Antibody Directed against Neisseria gonorrhoeae Lipooligosaccharide. Journal of immunology (Baltimore, Md : 1950) 196, 4576–4586.

Chen, A., and Seifert, H.S. (2013). Structure-function studies of the Neisseria gonorrhoeae major outer membrane porin. Infection and immunity 81, 4383–4391.

Chinery, L., Wahome, N., Moal, I., and Deane, C.M. (2023). Paragraph-antibody paratope prediction using graph neural networks with minimal feature vectors. Bioinformatics (Oxford, England) 39.

Cohen, M.S., and Cannon, J.G. (1999). Human experimentation with Neisseria gonorrhoeae: progress and goals. The Journal of infectious diseases 179 Suppl 2, S375–379.

De Goeij, B.E., Janmaat, M.L., Andringa, G., Kil, L., Van Kessel, B., Frerichs, K.A., Lingnau, A., Freidig, A., Mutis, T., Sasser, A.K., et al. (2019). Hexabody-CD38, a Novel CD38 Antibody with a Hexamerization Enhancing Mutation, Demonstrates Enhanced Complement-Dependent Cytotoxicity and Shows Potent Anti-Tumor Activity in Preclinical Models of Hematological Malignancies. Blood 134, 3106–3106.

de Jong, R.N., Beurskens, F.J., Verploegen, S., Strumane, K., van Kampen, M.D., Voorhorst, M., Horstman, W., Engelberts, P.J., Oostindie, S.C., Wang, G., et al. (2016). A Novel Platform for the Potentiation of Therapeutic Antibodies Based on Antigen-Dependent Formation of IgG Hexamers at the Cell Surface. PLoS biology 14, e1002344.

Donnarumma, D., Maestri, C., Giammarinaro, P.I., Capriotti, L., Bartolini, E., Veggi, D., Petracca, R., Scarselli, M., and Norais, N. (2018). Native State Organization of Outer Membrane Porins Unraveled by HDx-MS. Journal of proteome research 17, 1794–1800.

Donnelly, J., Medini, D., Boccadifuoco, G., Biolchi, A., Ward, J., Frasch, C., Moxon, E.R., Stella, M., Comanducci, M., Bambini, S., et al. (2010). Qualitative and quantitative assessment of meningococcal antigens to evaluate the potential strain coverage of protein-based vaccines. Proceedings of the National Academy of Sciences of the United States of America 107, 19490–19495.

Ferrari, G., Garaguso, I., Adu-Bobie, J., Doro, F., Taddei, A.R., Biolchi, A., Brunelli, B., Giuliani, M.M., Pizza, M., Norais, N., et al. (2006). Outer membrane vesicles from group B Neisseria meningitidis delta gna33 mutant: proteomic and immunological comparison with detergent-derived outer membrane vesicles. Proteomics 6, 1856–1866.

Findlow, J., Holland, A., Andrews, N., Weynants, V., Sotolongo, F., Balmer, P., Poolman, J., and Borrow, R. (2007). Comparison of phenotypically indistinguishable but geographically distinct Neisseria meningitidis Group B isolates in a serum bactericidal antibody assay. Clinical and vaccine immunology : CVI 14, 1451–1457.

Goire, N., Lahra, M.M., Chen, M., Donovan, B., Fairley, C.K., Guy, R., Kaldor, J., Regan, D., Ward, J., Nissen, M.D., et al. (2014). Molecular approaches to enhance surveillance of gonococcal antimicrobial resistance. Nature reviews Microbiology 12, 223–229.

Gottlieb, S.L., Ndowa, F., Hook, E.W., 3rd, Deal, C., Bachmann, L., Abu-Raddad, L., Chen, X.S., Jerse, A., Low, N., MacLennan, C.A., et al. (2020). Gonococcal vaccines: Public health value and preferred product characteristics; report of a WHO global stakeholder consultation, January 2019. Vaccine 38, 4362–4373.

Greenberg, L. (1975). Field trials of a gonococcal vaccine. The Journal of reproductive medicine 14, 34–36.

Gulati, S., Beurskens, F.J., de Kreuk, B.-J., Roza, M., Zheng, B., DeOliveira, R.B., Shaughnessy, J., Nowak, N.A., Taylor, R.P., Botto, M., et al. (2019a). Complement alone drives efficacy of a chimeric antigonococcal monoclonal antibody. PLoS biology 17, e3000323.

Gulati, S., Pennington, M.W., Czerwinski, A., Carter, D., Zheng, B., Nowak, N.A., DeOliveira, R.B., Shaughnessy, J., Reed, G.W., Ram, S., et al. (2019b). Preclinical Efficacy of a Lipooligosaccharide Peptide Mimic Candidate Gonococcal Vaccine. mBio 10.

Heckels, J.E., Virji, M., Zak, K., and Fletcher, J.N. (1987). Immunobiology of gonococcal outer membrane protein I. Antonie van Leeuwenhoek 53, 461–464.

Hill, S.A., Masters, T.L., and Wachter, J. (2016). Gonorrhea - an evolving disease of the new millennium. Microbial cell (Graz, Austria) 3, 371–389.

Hobbs, M.M., Sparling, P.F., Cohen, M.S., Shafer, W.M., Deal, C.D., and Jerse, A.E. (2011). Experimental Gonococcal Infection in Male Volunteers: Cumulative Experience with Neisseria gonorrhoeae Strains FA1090 and MS11mkC. Frontiers in microbiology 2, 123.

Holst, J., Oster, P., Arnold, R., Tatley, M.V., Næss, L.M., Aaberge, I.S., Galloway, Y., McNicholas, A., O’Hallahan, J., Rosenqvist, E., et al. (2013). Vaccines against meningococcal serogroup B disease containing outer membrane vesicles (OMV): lessons from past programs and implications for the future. Human vaccines & immunotherapeutics 9, 1241–1253.

Huang, J., Doria-Rose, N.A., Longo, N.S., Laub, L., Lin, C.L., Turk, E., Kang, B.H., Migueles, S.A., Bailer, R.T., Mascola, J.R., et al. (2013). Isolation of human monoclonal antibodies from peripheral blood B cells. Nature protocols 8, 1907–1915.

Humphries, H.E., Williams, J.N., Blackstone, R., Jolley, K.A., Yuen, H.M., Christodoulides, M., and Heckels, J.E. (2006). Multivalent liposome-based vaccines containing different serosubtypes of PorA protein induce cross-protective bactericidal immune responses against Neisseria meningitidis. Vaccine 24, 36–44.

Jansen, C., Kuipers, B., van der Biezen, J., de Cock, H., van der Ley, P., and Tommassen, J. (2000). Immunogenicity of in vitro folded outer membrane protein PorA of Neisseria meningitidis. FEMS immunology and medical microbiology 27, 227–233.

Jarvis, G.A., and Chang, T.L. (2012). Modulation of HIV transmission by Neisseria gonorrhoeae: molecular and immunological aspects. Current HIV research 10, 211–217.

Jennings, M.P., Srikhanta, Y.N., Moxon, E.R., Kramer, M., Poolman, J.T., Kuipers, B., and van der Ley, P. (1999). The genetic basis of the phase variation repertoire of lipopolysaccharide immunotypes in Neisseria meningitidis. Microbiology (Reading, England) 145 (Pt 11), 3013–3021.

Jerse, A.E. (1999). Experimental gonococcal genital tract infection and opacity protein expression in estradiol-treated mice. Infection and immunity 67, 5699–5708.

Joiner, K.A., Warren, K.A., Tam, M., and Frank, M.M. (1985). Monoclonal antibodies directed against gonococcal protein I vary in bactericidal activity. Journal of immunology (Baltimore, Md : 1950) 134, 3411-3419.

Jones, R.A., Jerse, A.E., and Tang, C.M. (2023). Gonococcal PorB: a multifaceted modulator of host immune responses. Trends in microbiology.

Jumper, J., Evans, R., Pritzel, A., Green, T., Figurnov, M., Ronneberger, O., Tunyasuvunakool, K., Bates, R., Žídek, A., Potapenko, A., et al. (2021). Highly accurate protein structure prediction with AlphaFold. Nature 596, 583–589.

Lenz, J.D., and Dillard, J.P. (2018). Pathogenesis of Neisseria gonorrhoeae and the Host Defense in Ascending Infections of Human Fallopian Tube. Frontiers in immunology 9, 2710.

Longtin, J., Dion, R., Simard, M., Betala Belinga, J.F., Longtin, Y., Lefebvre, B., Labbé, A.C., Deceuninck, G., and De Wals, P. (2017). Possible Impact of Wide-scale Vaccination Against Serogroup B Neisseria Meningitidis on Gonorrhea Incidence Rates in One Region of Quebec, Canada (Open Forum Infect Dis. 2017 Oct 4;4(Suppl 1):S734–5. doi: 10.1093/ofid/ofx180.002. eCollection 2017 Fall.).

Malito, E., Faleri, A., Lo Surdo, P., Veggi, D., Maruggi, G., Grassi, E., Cartocci, E., Bertoldi, I., Genovese, A., Santini, L., et al. (2013). Defining a protective epitope on factor H binding protein, a key meningococcal virulence factor and vaccine antigen. Proceedings of the National Academy of Sciences of the United States of America 110, 3304–3309.

Manca, B., Buffi, G., Magri, G., Del Vecchio, M., Taddei, A.R., Pezzicoli, A., and Giuliani, M. (2023). Functional characterization of the gonococcal polyphosphate pseudo-capsule. PLoS pathogens 19, e1011400.

Mandrell, R.E., Griffiss, J.M., and Macher, B.A. (1988). Lipooligosaccharides (LOS) of Neisseria gonorrhoeae and Neisseria meningitidis have components that are immunochemically similar to precursors of human blood group antigens. Carbohydrate sequence specificity of the mouse monoclonal antibodies that recognize crossreacting antigens on LOS and human erythrocytes. The Journal of experimental medicine 168, 107–126.

Marjuki, H., Topaz, N., Joseph, S.J., Gernert, K.M., Kersh, E.N., and Wang, X. (2019). Genetic Similarity of Gonococcal Homologs to Meningococcal Outer Membrane Proteins of Serogroup B Vaccine. mBio 10.

Martinón-Torres, F., Banzhoff, A., Azzari, C., De Wals, P., Marlow, R., Marshall, H., Pizza, M., Rappuoli, R., and Bekkat-Berkani, R. (2021). Recent advances in meningococcal B disease prevention: real-world evidence from 4CMenB vaccination. Journal of Infection 83, 17–26.

Marzoa, J., Sánchez, S., Ferreirós, C.M., and Criado, M.T. (2010). Identification of Neisseria meningitidis outer membrane vesicle complexes using 2-D high resolution clear native/SDS-PAGE. Journal of proteome research 9, 611–619.

Maurakis, S.A., and Cornelissen, C.N. (2022). Recent Progress Towards a Gonococcal Vaccine. Frontiers in cellular and infection microbiology 12, 881392.

McIntosh, E.D., Bröker, M., Wassil, J., Welsch, J.A., and Borrow, R. (2015). Serum bactericidal antibody assays - The role of complement in infection and immunity. Vaccine 33, 4414–4421.

McLeod Griffiss, J., Brandt, B.L., Saunders, N.B., and Zollinger, W. (2000). Structural relationships and sialylation among meningococcal L1, L8, and L3,7 lipooligosaccharide serotypes. The Journal of biological chemistry 275, 9716–9724.

Micoli, F., Ravenscroft, N., Cescutti, P., Stefanetti, G., Londero, S., Rondini, S., and Maclennan, C.A. (2014). Structural analysis of O-polysaccharide chains extracted from different Salmonella Typhimurium strains. Carbohydrate research 385, 1–8.

Młynarczyk-Bonikowska, B., Majewska, A., Malejczyk, M., Młynarczyk, G., and Majewski, S. (2020). Multiresistant Neisseria gonorrhoeae: a new threat in second decade of the XXI century. Medical microbiology and immunology 209, 95–108.

Mubaiwa, T.D., Semchenko, E.A., Hartley-Tassell, L.E., Day, C.J., Jennings, M.P., and Seib, K.L. (2017). The sweet side of the pathogenic Neisseria: the role of glycan interactions in colonisation and disease. Pathogens and disease 75.

O’Connor, E.T., Swanson, K.V., Cheng, H., Fluss, K., Griffiss, J.M., and Stein, D.C. (2008). Structural requirements for monoclonal antibody 2-1-L8 recognition of neisserial lipooligosaccharides. Hybridoma (2005) 27, 71–79.

Ochoa-Azze, R.F. (2018). Cross-protection induced by VA-MENGOC-BC® vaccine. Human vaccines & immunotherapeutics 14, 1064–1068.

Oostindie, S.C., van der Horst, H.J., Kil, L.P., Strumane, K., Overdijk, M.B., van den Brink, E.N., van den Brakel, J.H.N., Rademaker, H.J., van Kessel, B., van den Noort, J., et al. (2020). DuoHexaBody-CD37(®), a novel biparatopic CD37 antibody with enhanced Fc-mediated hexamerization as a potential therapy for B-cell malignancies. Blood cancer journal 10, 30.

Otsu, N. (1979). A Threshold Selection Method from Gray-Level Histograms. IEEE Transactions on Systems, Man, and Cybernetics 9, 62–66.

Paynter, J., Goodyear-Smith, F., Morgan, J., Saxton, P., Black, S., and Petousis-Harris, H. (2019). Effectiveness of a Group B Outer Membrane Vesicle Meningococcal Vaccine in Preventing Hospitalization from Gonorrhea in New Zealand: A Retrospective Cohort Study. Vaccines 7.

Pizza, M., Scarlato, V., Masignani, V., Giuliani, M.M., Aricò, B., Comanducci, M., Jennings, G.T., Baldi, L., Bartolini, E., Capecchi, B., et al. (2000). Identification of vaccine candidates against serogroup B meningococcus by whole-genome sequencing. Science (New York, NY) 287, 1816–1820.

Ram, S., Cullinane, M., Blom, A.M., Gulati, S., McQuillen, D.P., Monks, B.G., O’Connell, C., Boden, R., Elkins, C., Pangburn, M.K., et al. (2001). Binding of C4b-binding protein to porin: a molecular mechanism of serum resistance of Neisseria gonorrhoeae. The Journal of experimental medicine 193, 281–295.

Ram, S., Gulati, S., Lewis, L.A., Chakraborti, S., Zheng, B., DeOliveira, R.B., Reed, G.W., Cox, A.D., Li, J., St Michael, F., et al. (2018). A Novel Sialylation Site on Neisseria gonorrhoeae Lipooligosaccharide Links Heptose II Lactose Expression with Pathogenicity. Infection and immunity 86.

Ruffolo, J.A., Sulam, J., and Gray, J.J. (2022). Antibody structure prediction using interpretable deep learning. Patterns 3, 100406.

Russell, M.W., Jerse, A.E., and Gray-Owen, S.D. (2019). Progress Toward a Gonococcal Vaccine: The Way Forward. Frontiers in immunology 10, 2417.

Sacchi, C.T., Lemos, A.P.S., Brandt, M.E., Whitney, A.M., Melles, C.E.A., Solari, C.A., Frasch, C.E., and Mayer, L.W. (1998). Proposed Standardization of Neisseria meningitidis PorA Variable-Region Typing Nomenclature. 5, 845–855.

Semchenko, E.A., Tan, A., Borrow, R., and Seib, K.L. (2019). The Serogroup B Meningococcal Vaccine Bexsero Elicits Antibodies to Neisseria gonorrhoeae. Clinical infectious diseases : an official publication of the Infectious Diseases Society of America 69, 1101–1111.

Snape, M.D., Saroey, P., John, T.M., Robinson, H., Kelly, S., Gossger, N., Yu, L.M., Wang, H., Toneatto, D., Dull, P.M., et al. (2013). Persistence of bactericidal antibodies following early infant vaccination with a serogroup B meningococcal vaccine and immunogenicity of a preschool booster dose. CMAJ : Canadian Medical Association journal = journal de l’Association medicale canadienne 185, E715–724.

St Cyr, S., Barbee, L., Workowski, K.A., Bachmann, L.H., Pham, C., Schlanger, K., Torrone, E., Weinstock, H., Kersh, E.N., and Thorpe, P. (2020). Update to CDC’s Treatment Guidelines for Gonococcal Infection, 2020. MMWR Morbidity and mortality weekly report 69, 1911–1916.

Stazzoni, S., Troisi, M., Abbiento, V., Sala, C., Andreano, E., and Rappuoli, R. (2023). High-throughput bactericidal assays for monoclonal antibody screening against antimicrobial resistant Neisseria gonorrhoeae. Frontiers in microbiology 14, 1243427.

Tani, C., Stella, M., Donnarumma, D., Biagini, M., Parente, P., Vadi, A., Magagnoli, C., Costantino, P., Rigat, F., and Norais, N. (2014). Quantification by LC-MS(E) of outer membrane vesicle proteins of the Bexsero® vaccine. Vaccine 32, 1273–1279.

Tinsley, C.R., and Nassif, X. (1996). Analysis of the genetic differences between Neisseria meningitidis and Neisseria gonorrhoeae: two closely related bacteria expressing two different pathogenicities. Proceedings of the National Academy of Sciences of the United States of America 93, 11109–11114.

Tondella, M.L.C., Popovic, T., Rosenstein, N.E., Lake, D.B., Carlone, G.M., Mayer, L.W., and Perkins, B.A. (2000). Distribution of Neisseria meningitidisSerogroup B Serosubtypes and Serotypes Circulating in the United States. 38, 3323–3328.

Tramont, E.C. (1989). Gonococcal vaccines. Clinical microbiology reviews 2 *Suppl*, S74–77.

Tsai, C.M., and Civin, C.I. (1991). Eight lipooligosaccharides of Neisseria meningitidis react with a monoclonal antibody which binds lacto-N-neotetraose (Gal beta 1-4GlcNAc beta 1-3Gal beta 1-4Glc). Infection and immunity 59, 3604–3609.

Unemo, M., Lahra, M.M., Escher, M., Eremin, S., Cole, M.J., Galarza, P., Ndowa, F., Martin, I., Dillon, J.R., Galas, M., et al. (2021). WHO global antimicrobial resistance surveillance for Neisseria gonorrhoeae 2017-18: a retrospective observational study. The Lancet Microbe 2, e627–e636.

Unemo, M., Norlén, O., and Fredlund, H. (2005). The porA pseudogene of Neisseria gonorrhoeae--low level of genetic polymorphism and a few, mainly identical, inactivating mutations. APMIS : acta pathologica, microbiologica, et immunologica Scandinavica 113, 410–419.

Unemo, M., and Shafer, W.M. (2014). Antimicrobial resistance in Neisseria gonorrhoeae in the 21st century: past, evolution, and future. Clinical microbiology reviews 27, 587–613.

van Zundert, G.C.P., Rodrigues, J., Trellet, M., Schmitz, C., Kastritis, P.L., Karaca, E., Melquiond, A.S.J., van Dijk, M., de Vries, S.J., and Bonvin, A. (2016). The HADDOCK2.2 Web Server: User-Friendly Integrative Modeling of Biomolecular Complexes. Journal of molecular biology 428, 720–725.

Vipond, C., Suker, J., Jones, C., Tang, C., Feavers, I.M., and Wheeler, J.X. (2006). Proteomic analysis of a meningococcal outer membrane vesicle vaccine prepared from the group B strain NZ98/254. Proteomics 6, 3400–3413.

Viviani, V., Fantoni, A., Tomei, S., Marchi, S., Luzzi, E., Bodini, M., Muzzi, A., Giuliani, M.M., Maione, D., Derrick, J.P., et al. (2023). OpcA and PorB are novel bactericidal antigens of the 4CMenB vaccine in mice and humans. npj Vaccines 8, 54.

Wang, B., Giles, L., Andraweera, P., McMillan, M., Almond, S., Beazley, R., Mitchell, J., Lally, N., Ahoure, M., Denehy, E., et al. (2022). Effectiveness and impact of the 4CMenB vaccine against invasive serogroup B meningococcal disease and gonorrhoea in an infant, child, and adolescent programme: an observational cohort and case-control study. The Lancet Infectious Diseases 22, 1011–1020.

Westphal, O.a.J. K. (1965). Bacterial Lipopolysaccharides Extraction with Phenol-Water and Further Applications of the Procedure. Methods in Carbohydrate Chemistry 5, 83–91.

Whelan, J., Kløvstad, H., Haugen, I.L., Holle, M.R., and Storsaeter, J. (2016). Ecologic Study of Meningococcal B Vaccine and Neisseria gonorrhoeae Infection, Norway. Emerging infectious diseases 22, 1137–1139.

Yamasaki, R., Koshino, H., Kurono, S., Nishinaka, Y., McQuillen, D.P., Kume, A., Gulati, S., and Rice, P.A. (1999). Structural and immunochemical characterization of a Neisseria gonorrhoeae epitope defined by a monoclonal antibody 2C7; the antibody recognizes a conserved epitope on specific lipo-oligosaccharides in spite of the presence of human carbohydrate epitopes. The Journal of biological chemistry 274, 36550–36558.

